# *bak1-5* mutation uncouples tryptophan-dependent and independent postinvasive immune pathways triggered in Arabidopsis by multiple fungal pathogens

**DOI:** 10.1101/2020.04.26.052480

**Authors:** Ayumi Kosaka, Marta Pastorczyk, Mariola Piślewska-Bednarek, Takumi Nishiuchi, Haruka Suemoto, Atsushi Ishikawa, Henning Frerigmann, Masanori Kaido, Kazuyuki Mise, Paweł Bednarek, Yoshitaka Takano

## Abstract

Robust nonhost resistance of *Arabidopsis thaliana* against the nonadapted hemibiotrophic fungus *Colletotrichum tropicale* requires PEN2-dependent preinvasive and CYP71A12/CYP71A13-dependent postinvasive resistance, which both rely on tryptophan (Trp) metabolism. Here we report that CYP71A12 and CYP71A13 are critical for Arabidopsis’ postinvasive resistance toward both the necrotrophic *Alternaria brassicicola* and the adapted hemibiotrophic *C. higginsianum* fungi. Metabolite analyses suggest that the production of indole-3-carboxylic acid derivatives (ICAs) and camalexin is induced upon pathogen invasion, while phenotypic comparison of *cyp79B2 cyp79B3* and *pen2 cyp71A12 cyp71A13* plants indicates that the contribution of ICAs to postinvasive resistance is dose-dependent. We also found that the disruption of intact pattern recognition receptor complex caused by *bak1–5* mutation significantly reduced postinvasive resistance against *C. tropicale* and *A. brassicicola*, indicating that pattern recognition commonly contributes to this second defense-layer against pathogens with distinct infection strategies. However, the *bak1–5* mutation had no detectable effects on Trp-metabolite accumulation triggered by pathogen invasion. Together with this, further comparative gene expression analyses suggested that pathogen invasion in Arabidopsis activates (i) *bak1–5* insensitive Trp-metabolism that leads to antimicrobial secondary metabolites, and (ii) a *bak1–5* sensitive immune pathway that activates the expression of antimicrobial proteins.

## INTRODUCTION

The resistance of an entire plant species against all isolates of particular pathogens is called nonhost resistance [1]. *Arabidopsis thaliana* exhibits nonhost resistance against a nonadapted powdery mildew pathogen *Blumeria graminis* f. sp. *hordei* called *Bgh*. The four *PENETRATION* genes, *PEN1, PEN2, PEN3* and *PEN4*, have been reported in Arabidopsis to control entry of *Bgh* through activation of vesicle secretion, biosynthesis of putatively antifungal metabolites, and their transport toward pathogen invasion sites, respectively [2,3,4,5,6,7,8,9,10].

*Colletotrichum tropicale* (hereafter *Ctro*), formally called *C. gleosporioides*, is a hemibiotrophic fungal pathogen that causes anthracnose on its host mulberry; however, it is not able to infect the nonhost Arabidopsis. Entry control by Arabidopsis against *Ctro* involves *PEN2* and *PEN3* [11,12], whereas *PEN1* is not essential for this unlike the case of *Bgh* [13]. Nonhost resistance toward *Ctro* also needs *EDR1 (ENHANCED DISEASE RESISTANCE 1*) [14], whereas the *edr1* mutants exhibit enhanced resistance toward the adapted powdery mildew *Golovinomyces cichoracearum* (formerly named *Erysiphe cichoracearum*) [15], suggesting the presence of diverse plant strategies for controlling the entry attempts of pathogens showing distinct infection modes.

Importantly, even the *pen2 edr1* mutant is still not fully susceptible to *Ctro* [16], because of strong postinvasive resistance activated once *Ctro* enters epidermal cells of this mutant, which is defective in the preinvasive resistance that controls pathogen entry. We reported previously that Arabidopsis *cyp79B2 cyp79B3* double mutant is fully susceptible to the nonadapted pathogen *Ctro* [16]. CYP79B2 and CYP79B3 are key enzymes for the biosynthesis of tryptophan (Trp)-derived antimicrobial metabolites. CYP79B2/CYP79B3 convert Trp into indole-3-acetaldoxime (IAOx) [17], and this precursor is then converted into several compounds for antimicrobial immunity, such as PEN2 substrates indole-glucosinolates (IGs), PAD3 (PHYTOALEXIN-DEFICIENT3)-dependent camalexin, and 4-hydroxy-ICN (4-OH-ICN), whose biosynthesis requires CYP82C2 [18,19,20]. The *cyp79B2 cyp79B3* mutant is defective in preinvasive resistance against *Ctro* because CYP79B2 and CYP79B3 are indispensable for production of IGs, which are substrates metabolized by PEN2 to (unidentified) compounds with presumed antifungal activity [5]. However, contrary with the *pen2* mutant, the *cyp79B2 cyp79B3* plants were also defective in postinvasive resistance to *Ctro* [16]. We have shown that the *pen2 pad3* mutant is partially defective in postinvasive resistance to *Ctro*, indicating that the Arabidopsis phytoalexin, camalexin, is a critical factor for this. Importantly, the *cyp79B2 cyp79B3* mutant plants exhibit a more severe defect in this second-defense layer than the *pen2 pad3* plants [16]. This enhanced susceptibility in the *cyp79B2 cyp79B3* compared with the *pen2 pad3* plants was also reported for the interactions with *Phytophthora brassicae* and *Plectosphaerella cucumerina* [21,22]. These findings strongly suggested the presence of additional Trp-derived secondary metabolites that might be crucial for postinvasive resistance against fungal pathogens.

In addition to serving as a precursor of IGs and camalexin, IAOx can be also converted to indole-3-carboxylic acid and its derivatives (ICAs). We have shown recently that CYP71A12 but not CYP71A13 has an important contribution to the accumulation of ICAs in leaves upon both *Ctro* and *P. cucumerina* inoculation [23]. On the other hand, loss of CYP71A13 reduced camalexin accumulation in leaves upon infection by multiple pathogens [23,24], whereas a single loss of CYP71A12 did not reduce camalexin accumulation upon *Ctro* and *P. cucumerina* infection [23]. These findings suggest distinct roles of these two homologous P450 monooxygenases in the responses toward pathogen infection. Importantly, the *pen2 cyp71A12* double mutant exhibits a partial reduction in postinvasive resistance to *Ctro* [23], which was similar to that of the *pen2 pad3* plants. Furthermore, the *pen2 cyp71A12 cyp71A13* plants exhibited enhanced susceptibility compared with the *pen2 pad3* and *pen2 cyp71A12* plants. These findings suggest that CYP71A12-dependent synthesis of ICAs as well as camalexin synthesis is critical for postinvasive resistance to *Ctro*.

It has been reported that in addition to the above mentioned pests, camalexin is critical for the immunity of Arabidopsis to additional filamentous pathogens including necrotrophic fungus *Alternaria brassicicola* (hereafter called *Ab*). This indicated a common role of camalexin in the postinvasive antifungal defense in this plant species [16,22,25,26]. However, it remains unclear whether CYP71A12 and ICAs are commonly involved in the postinvasive resistance to fungal pathogens other than *Ctro*.

Here, we investigated whether CYP71A12-dependent synthesis of ICAs might be involved in postinvasive resistance to a necrotrophic fungal pathogen *Ab* that infects Brassicaceae plants. We found that invasion by *Ab* triggers the CYP71A12-dependent accumulation of ICAs as well as camalexin, and that postinvasive resistance to *Ab* needs both CYP71A12 and PAD3. We also reveal a similar function of these compounds in the postinvasive resistance to the adapted hemibiotrophic fungus *Colletotrichum higginsianum* (hereafter called *Ch*), suggesting common roles of CYP71A12 and ICAs for this invasion-triggered defense against diverse fungal pathogens with distinct infection modes.

BAK1 (BRASSINOSTEROID INSENSITIVE 1-ASSOCIATED RECEPTOR KINASE 1) is known to act as a coreceptor with multiple pattern recognition-receptor (PRRs), including FLS2 and EFR, via ligand-induced heteromerization [27,28,29,30]. BAK1 was initially identified as a positive regulator of the brassinosteroid response (BR) [31,32]. Correspondingly, the *bak1* null mutants such as *bak1-4* mutant are not only defective in FLS2 and EFR-dependent immune responses, but also hyposensitive to BR. In contrast to the null alleles, the *bak1–5* allele is impaired in pathogen-associated molecular pattern (PAMP)-triggered immunity, but not in BR signaling [33]. We investigated function of BAK1 in resistance and ICA formation and found that the *bak1* mutations, especially *bak1–5*, reduce postinvasive resistance to *Ab*, revealing the involvement of a PRR system in the recognition of pathogen invasion for the activation of defense. Contrarily, we reveal that the *bak1–5* mutation has no negative impact on the invasion-triggered activation of camalexin or ICA biosynthesis.

## RESULTS

### CYP71A12 contributes to the immunity of Arabidopsis against the necrotrophic pathogen *Alternaria brassicicola* independently of CYP71A13 and PAD3

*Ab* is a necrotrophic fungal pathogen that infects several Brassicaceae spp., including cabbage and canola but is restricted within limited lesions on the wild-type (WT) of *A. thaliana*, Col-0 [14,25,34]. To investigate the roles of Trp metabolism in the immunity of Arabidopsis against *Ab*, we inoculated the Ryo-1 strain of *Ab* onto the leaves of a series of Arabidopsis mutants related to Trp metabolism: *pen2, pen2 pad3, pen2 cyp82C2, pen2 cyp71A12, pen2 cyp71A12 cyp71A13*, and *cyp79B2 cyp79B3*. At 4 days postinoculation (dpi), lesion development was evaluated. Lesion development in the *pen2* mutant was not significantly different from that in the WT Col-0 leaves (Fig. 1A, B), suggesting that opposite with the impact on several filamentous pathogens, PEN2 has no detectable contribution to the immunity of Arabidopsis against *Ab* [3,11]. Interestingly, our assays revealed that both the *cyp71A12* and *pen2 cyp71A12* mutants showed enhanced susceptibility to *Ab*, as compared with WT and *pen2* plants, indicating contribution of CYP71A12 to the immunity towards this fungus (Fig. 1; Supplementary Fig. S1).

**Figure 1.**
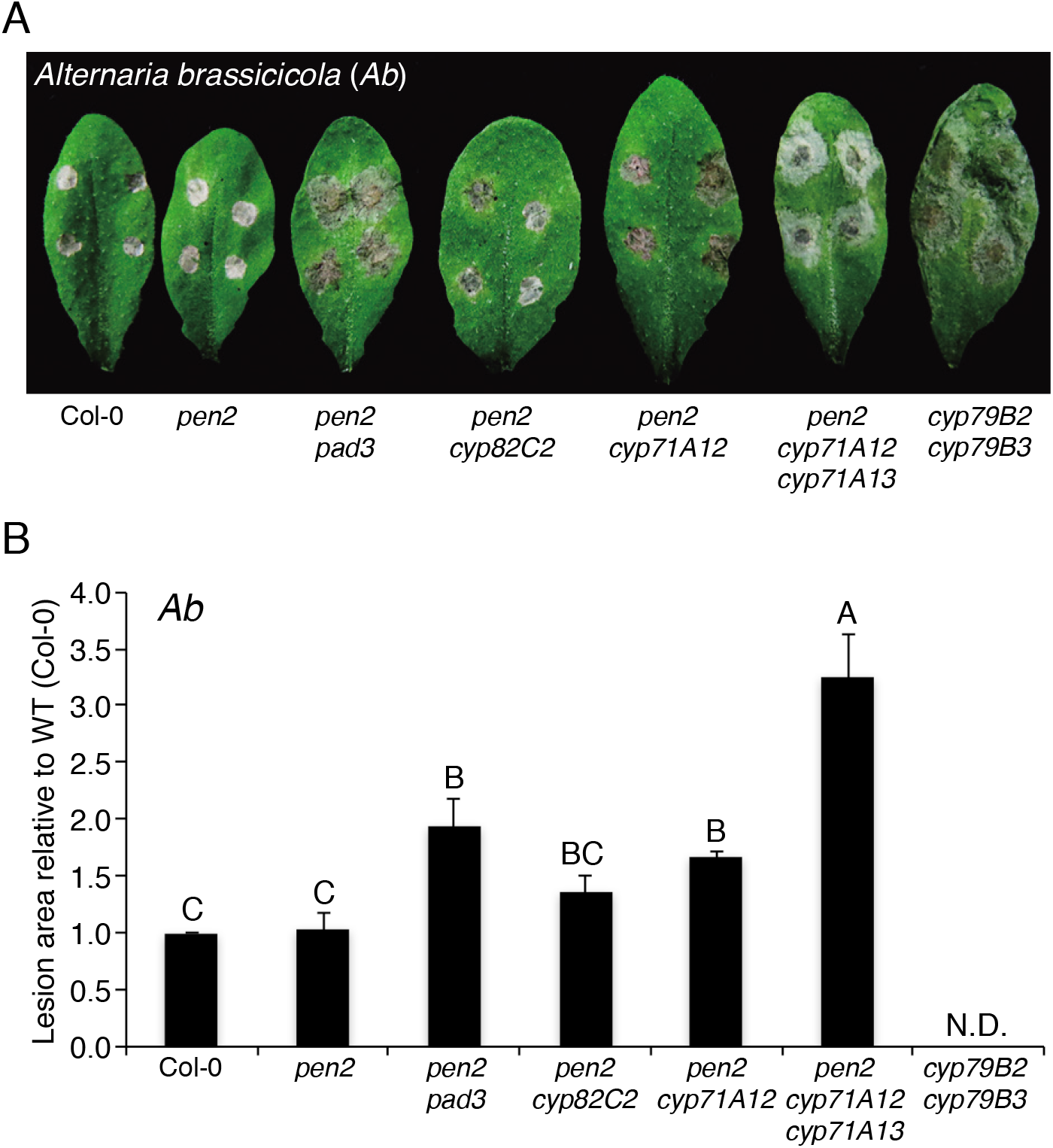
*PAD3* and *CYP71A12* are involved in the immunity of Arabidopsis against the necrotrophic pathogen *Alternaria brassicicola* (*Ab*). **(A)** Lesion development caused by *Ab* on Arabidopsis mutant plants with defects in Trp metabolism pathways. Conidial suspensions (1 × 10^5^ conidia/mL) of *Ab* were drop-inoculated onto mature leaves of 4–5-week-old plants. The photograph was taken at 4 days postinoculation (dpi). **(B)** Quantification of lesion development. Conidial suspensions of *Ab* were drop-inoculated onto tested plants. At 4 dpi, lesion areas were measured and the relative values to Col-0 (WT plants) were calculated. Means and standard deviations (SDs) were calculated from three independent experiments. The statistical significance of differences between means was determined by Tukey’s honestly significant difference (HSD) test. Means not sharing the same letter are significantly different (*P* < 0.05).

An additional mutation in *CYP71A13* further increased lesion development in the respective double (*cyp71A12 cyp71A13*) and triple (*pen2 cyp71A12 cyp71A13*) mutants. We also found that both the *pad3* and *pen2 pad3* mutants have increased susceptibility to the *Ab* Ryo-1 compared with the WT plant (Fig. 1A, B; Supplementary Fig. S1), indicating the importance of PAD3-dependent camalexin synthesis, consistent with previous reports on *pad3* susceptibility toward different *Ab* strains [25,34]. Opposite with *PAD3* and *CYP71A12/A13 cyp82C2* mutation did not cause significant changes in *Ab* development suggesting that *CYP82C2* together with 4-OH-ICN do not contribute to the immunity toward this pathogen (Fig. 1A, B; Supplementary Fig. S1). We were unable to perform proper quantitative analysis of lesion development in the *cyp79B2 cyp79B3* mutant at 4 dpi because the lesions had already merged. However, quantitative analysis at 3 dpi revealed that the *cyp79B2 cyp79B3* mutant was the most susceptible to *Ab* among all tested genotypes (Supplementary Fig. S2).

### Biosynthesis of ICAs and camalexin occurs during postinvasive resistance toward *Ab*

Pastorczyk, M. *et al*. [23] reported that (i) the expression of *CYP71A12* and *PAD3* is strongly induced in the *pen2* mutant upon the inoculation of the non-adapted pathogen *Ctro* and (ii) *CYP71A12* and *PAD3* are involved in postinvasive resistance against *Ctro*. To assess whether *CYP71A12* and *PAD3* are expressed during postinvasive resistance against *Ab*, similar as against *Ctro*, we investigated the expression pattern of these genes at several time points after *Ab* inoculation (at 4, 12, 24, and 48 h postinoculation, hpi). To enable precise comparison of current results with our findings on postinvasive resistance against *Ctro* [23], we decided to use mutants in the *pen2* background in our study with *Ab*, although the lesion development assay revealed that *PEN2* is not essential for immunity toward this fungi (Fig. 1A, B; Supplementary Fig. S1). We found that the expression of *CYP71A12* and *PAD3* started to be induced at 12 hpi, and the corresponding expression levels were even stronger elevated at later time points (Fig. 2A). By contrast, we did not detect any induction at 4 hpi (Fig. 2A). In parallel, we also investigated the temporal infection behavior of *Ab* in Arabidopsis. We found that conidia of *Ab* had already germinated at 4 hpi, however, we did not detect any host invasion at this time (Fig. 2B, C). The fungus started to invade the plants at 12 hpi and the invasion ratio became elevated at later time points (Fig. 2B, C). These findings indicate a link between *CYP71A12* and *PAD3* induction and the initiation of host invasion in the *Ab*-Arabidopsis interactions, strongly suggesting that the expressions of *CYP71A12* and *PAD3* are triggered by *Ab* invasion. Furthermore, we also revealed that simultaneous loss of both *CYP71A12* and *CYP71A13* produced no detectable effects on the preinvasive resistance against *Ab* (Fig. 2D), suggesting that *CYP71A12* and *CYP71A13* are involved in the postinvasive resistance against this fungus. We also found that *PEN2* is dispensable for the preinvasive resistance against *Ab* (Fig. 2D), in contrast to *Ctro* [11].

**Figure 2.**
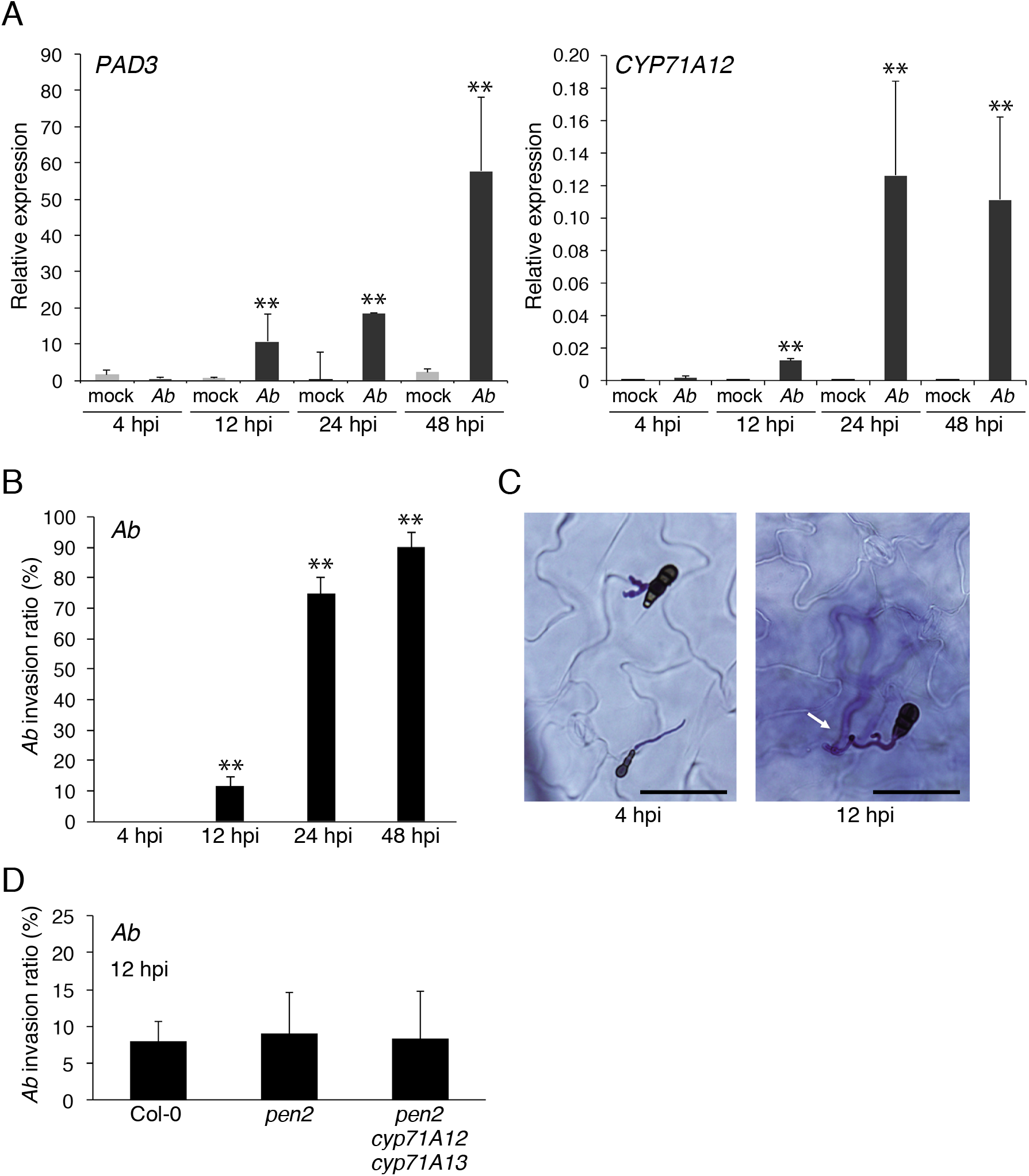
The invasion of *Ab* activates the expression of *PAD3* and *CYP71A12*. **(A)** Expression *PAD3* and *CYP71A12* following *Ab* inoculation. Conidial suspensions (5 × 10^5^ conidia/mL) of *Ab* were spray-inoculated onto 4–5-week-old *pen2* plants and kept at 100% humidity. The samples were collected at 4, 12, 24, and 48 hpi. Each gene transcript was quantified by quantitative polymerase chain reaction (qPCR) using the gene-specific primers listed in Supplementary Table S2. Values were normalized to the expression level of *UBC21*. The means and SDs were calculated from three independent experiments. Statistical comparisons between mock and *Ab* treated samples were conducted using two-tailed Student’s *t* tests (\*\**P* < 0.01). **(B)** Quantitative analysis of the *Ab* invasion ratio among *pen2* plants. Conidial suspensions (1 × 10^5^ conidia/mL) of *Ab* were drop-inoculated onto *pen2* plants and kept at 100% humidity. The inoculated leaves were collected at 4, 12, 24, and 48 hpi, and then subjected to a trypan blue viability staining assay. The presence or absence of invasive hyphae from at least 50 germinating conidia were counted in each experiment. The means and SDs were calculated from three independent experiments. Statistical comparisons of the *Ab* invasion ratios at 12, 24, and 48 hpi against that at 4 hpi were conducted using two-tailed Student’s *t* tests (\*\**P* < 0.01). **(C)** Light microscopy observations. At 4 hpi, most of the *Ab* conidia had germinated and started to elongate hyphae, but had not developed invasive hyphae. At 12 hpi, some of the germinating conidia developed invasive hyphae (arrow) inside the plants. Bars = 50 μm. **(D)** *PEN2, CYP71A12*, and *CYP71A13* are dispensable for preinvasive resistance against *Ab*. Aliquots of 5 μL of *Ab* conidial suspension were drop-inoculated onto leaves of 4–5-week-old plants. At 12 hpi, the inoculated leaves were collected and stained with trypan blue, and then invasive hyphae were observed by light microscopy. At least 50 germinating conidia were counted in each experiment. The means and SDs were calculated from three independent experiments. Statistical analysis using two-tailed Student’s *t* tests showed no significant differences among the genotypes.

We then assessed whether our observations made at the gene expression level could be supported with metabolite profiles. We found that the pathogen-induced accumulation of two ICAs, glucoside of 6-hydroxy-indole-3-carboxylic acid and glucose ester of indole-3-carboxylic acid, was detected at 24 hpi, but not at 4 hpi with *Ab* (Fig. 3A; Supplementary Fig. S3), which matched strongly with the gene expression data (Fig. 2A). Similar results were also obtained during the analysis of camalexin accumulation (Fig. 3A). Therefore, we conclude that the synthesis of ICAs and camalexin is triggered by *Ab* invasion.

**Figure 3.**
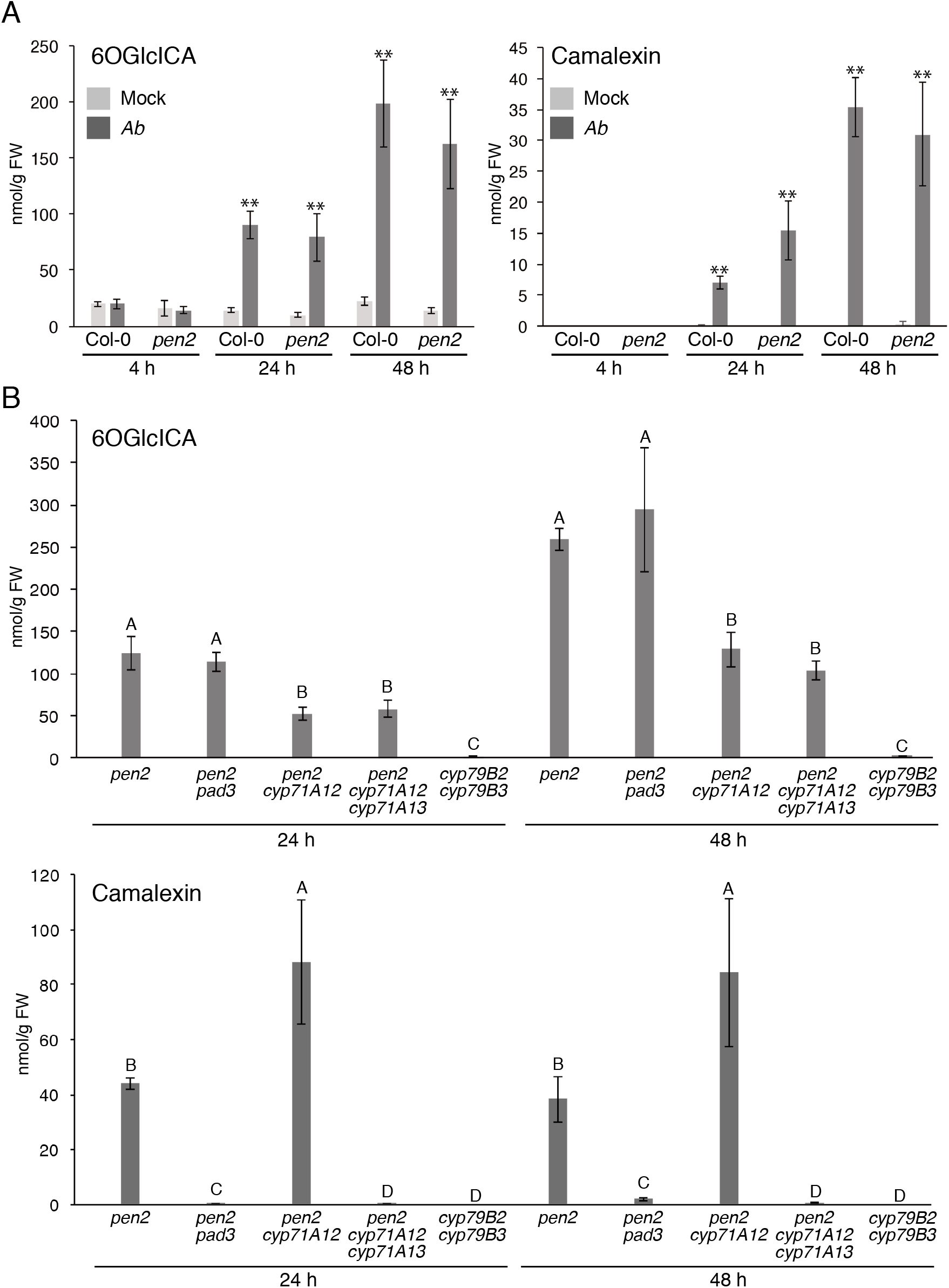
Invasion by *Ab* induced CYP7lAl2-dependent biosynthesis of 6-hydroxy-ICA and PAD3-dependent biosynthesis of camalexin. **(A)** The accumulation of 6-hydroxy-ICA (6OGlcICA) and camalexin in Arabidopsis plants inoculated with *Ab*. Conidial suspensions (5 × 10^5^ conidia/mL) of *Ab* were spray-inoculated onto Col-0 (WT) and *pen2* plants. As a control, water was sprayed as a mock treatment. The means of metabolites (nmol/g fresh weight, FW) and SDs from four biological replicates are shown in the graph. The statistical analysis between mock and *Ab* treated samples was conducted using twotailed Student’s *t* tests (\**P* < 0.05; \*\**P* < 0.01). **(B)** *CYP71A12* is required for the *Ab*-triggered accumulation of 6OG1cICA, whereas *PAD3* and *CYP71A13* are required for the accumulation of camalexin. Conidial suspensions (5 × 10^5^ conidia/mL) of *Ab* were spray-inoculated onto tested mutant lines. The statistical significance of differences between means was determined by Tukey’s honestly significant difference (HSD) test. Means not sharing the same letter are significantly different (*P* < 0.05).

### Reduced accumulation of ICAs correlates with the breakdown of the postinvasive resistance toward *Ab*

Our assays revealed that both the *cyp71A12* and *pen2 cyp71A12* mutants showed enhanced susceptibility to *Ab*, suggesting the importance of CYP71A12-dependent production of ICAs (Fig. 1A, B; Supplementary Fig. S1). Furthermore, we found that the loss of *CYP71A13* additionally reduced the immune response to *Ab* in both the *cyp71A12* and *pen2 cyp71A12* mutants (Fig. 1A, B; Supplementary Fig. S1). We have reported recently that the accumulation of ICAs triggered by inoculation with *P. cucumerina* is reduced in the *cyp71A12*, but not in the *cyp71A13* mutant [23]. By contrast, the *P. cucumerina*-triggered accumulation of camalexin was reduced in the *cyp71A13*, but not in the single *cyp71A12* mutant [23]. Therefore, we assumed that the CYP71A12-dependent production of ICAs contributes to the immunity of Arabidopsis to *Ab* independently of CYP71A13-dependent camalexin production, which is also consistent with the finding that the phenotype of the *pen2 pad3* mutant was less severe than that of the *pen2 cyp71A12 cyp71A13* mutant after *Ab* inoculation (Fig. 1A, B).

To validate this hypothesis, we investigated the Trp-related metabolite profiles of the aforementioned mutants: *pen2*, *pen2 pad3*, *pen2 cyp71A12*, *pen2 cyp71A12 cyp71A13*, and *cyp79B2 cyp79B3*. Obtained results revealed that the simultaneous loss of *CYP79B2* and *CYP79B3* completely abolished the *Ab*-triggered accumulation of ICAs and camalexin, whereas the loss of *PAD3* canceled camalexin accumulation, but had no effects on the accumulation of ICAs (Fig. 3B; Supplementary Fig. S4). In the *pen2 cyp71A12* mutant, the *Ab*-triggered accumulation of ICAs was reduced significantly compared with the *pen2* mutant, opposite with camalexin accumulation that was rather increased compared with *pen2* (Fig. 3B; Supplementary Fig. S4). These results further strengthen the idea that the accumulation of ICAs triggered by the *Ab* invasion is critical for the postinvasive resistance of Arabidopsis against this necrotrophic fungal pathogen. In the *pen2 cyp71A12 cyp71A13* mutant, the ICA levels that accumulated upon *Ab* invasion were similar to those observed in the *pen2 cyp71A12* mutant, but camalexin accumulation in the triple mutant was completely diminished, in contrast to *pen2 cyp71A12* (Fig. 3B; Supplementary Fig. S4), further supporting the importance of CYP71A13 for the pathogen-induced accumulation of camalexin, but not ICAs.

It is noteworthy that the leaves of *pen2 cyp71A12 cyp71A13* mutant still accumulated clearly detectable amounts of ICAs upon *Ab* inoculation, whereas camalexin in this triple mutant was under the detection limit (Fig. 3B; Supplementary Fig. S4). As mentioned above, the *cyp79B2 cyp79B3* mutant exhibited a more severe phenotype to *Ab* inoculation compared with the *pen2 cyp71A12 cyp71A13* mutant (Fig. 1A, B; Supplementary Fig. S2). The *cyp79B2 cyp79B3* mutant is entirely defective in the production of not only camalexin, but also ICAs (Fig. 3B; Supplementary Fig. S4). These data suggested that the different levels of susceptibility to *Ab* between *pen2 cyp71A12 cyp71A13* and *cyp79B2 cyp79B3* is likely caused by the differential accumulation of ICAs in these two mutants. Consistent with this idea, the *pen2 cyp71A12* mutant was more susceptible to *Ab* than the *pen2* mutant (Fig. 1) additionally supporting the correlation between the reduced levels of ICAs and the breakdown of the postinvasive resistance toward *Ab*.

In addition to ICAs and camalexin, CYP79B2/CYP89B3 enzymes are essential for biosynthesis of other Trp-derived metabolites including IGs, [23,35]. The *pen2 cyp71A12 cyp71A13* mutant lacks the PEN2-dependent IG-hydrolysis products, but retains the ability to produce IGs, which can be activated by another enzyme. It has been reported that IG biosynthesis in Arabidopsis is controlled by the transcription factors MYB34, MYB51, and MYB122, whereas these factors are dispensable for the pathogen triggered biosynthesis of camalexin and ICAs [35]. Thus, to assess the possibility that the different susceptibility to *Ab* between *pen2 cyp71A12 cyp71A13* and *cyp79B2 cyp79B3* mutants might be linked to IG biosynthesis, we compared phenotypes of *pen2 cyp71A12 cyp71A13* and *cyp71A12 cyp71A13 myb34 myb51 myb122* lines following *Ab* inoculation. Lesion development in the *cyp71A12 cyp71A13 myb34 myb51 myb122* mutant leaves was similar to that in the *pen2 cyp71A12 cyp71A13* mutant, suggesting that PEN2-independent IG hydrolysis is likely not involved in the postinvasive resistance against *Ab* (Supplementary Fig. S5). Thus, the different susceptibility between the *pen2 cyp71A12 cyp71A13* mutant and the *cyp79B2 cyp79B3* mutant is not linked to IG-deficiency. Together with the involvement of CYP71A12-dependent ICAs synthesis in the postinvasive resistance of Arabidopsis against *Ab*, we consider that the remaining ICAs in *pen2 cyp71A12 cyp71A13* are still effective against *Ab*, i.e., the ICAs contribute to the postinvasive resistance of Arabidopsis in a dose-5dependent manner.

### ICAs function commonly in the postinvasive resistance of Arabidopsis against multiple fungal pathogens with different infection modes

We further investigated the possible contributions of ICAs to the immune response of Arabidopsis against the hemibiotrophic fungus *C. higginsianum* (*Ch*), which is adapted to this plant species. We inoculated conidial suspensions of *Ch* to the aforementioned Arabidopsis mutants with altered Trp metabolism. As a result, *Ch*-triggered lesion development in the *pen2* mutant was similar to that in the WT Col-0 (Fig. 4A), whereas *PEN2* was reported to be involved in preinvasive resistance against *Ch* to some degree [14]. Compared with *pen2*, we found increased lesion development in both the *pen2 pad3* and *pen2 cyp71A12* mutants (Fig. 4A). Furthermore, we found an additive effect in *pen2 cyp71A12 cyp71A13* compared with *pen2 pad3* and *pen2 cyp71A12* (Fig. 4A). These tendencies are clearly consistent with phenotypes observed for both *Ab* (Fig. 1) and *Ctro* [23]. We also found that the *pen2 cyp71A12 cyp71A13* plants had no obvious defects in preinvasive resistance against *Ch* (Fig. 4B). Therefore, we conclude that ICAs and camalexin function commonly in the postinvasive resistance of Arabidopsis against multiple fungal pathogens: the necrotrophic fungus *Ab*, and the two hemibiotrophic fungi, *Ch* and *Ctro*. By contrast, PEN2-dependent preinvasive resistance is critical for *Ctro* and is partially effective to *Ch* [11,14], but is dispensable for *Ab* (Fig. 2D). Therefore, we assume that Arabidopsis has evolved to use conserved and common Trp-metabolism based mechanisms of postinvasive resistance against a broad range of fungal pathogens, whereas it has developed distinct mechanisms for controlling entry of these pathogens.

**Figure 4.**
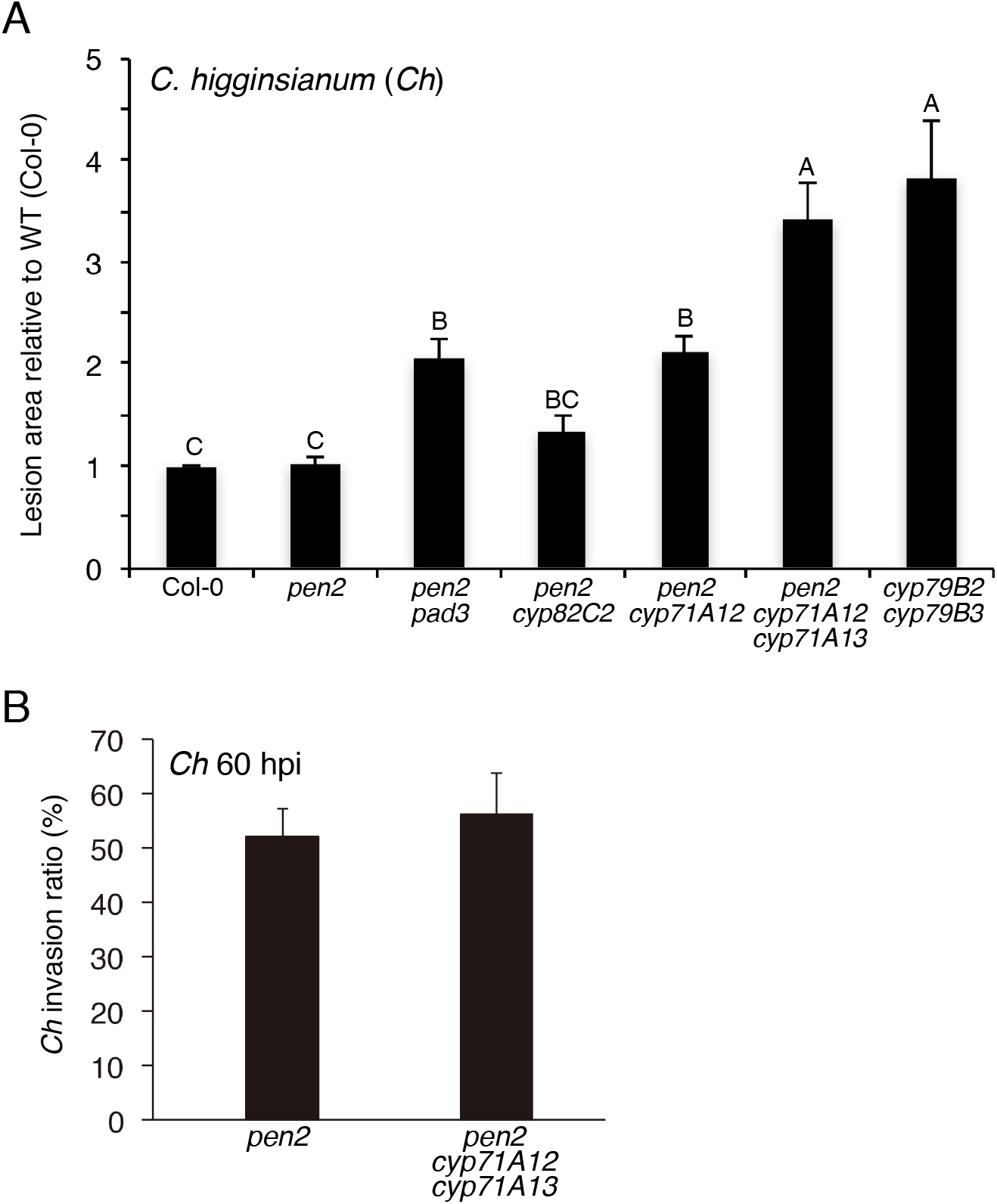
ICAs and camalexin are involved in the immunity of Arabidopsis against a hemibiotrophic pathogen *Colletotrichum higginsianum* (*Ch*). **(A)** Quantitative analysis of lesion development produced by *Ch*. Conidial suspensions of *Ch* (1 × 10^5^ conidia/mL) were drop-inoculated onto mature leaves of 4–5-week-old plants of the tested Arabidopsis lines. At 4 dpi, lesion development was measured. The means and SDs were calculated from three independent experiments. The statistical significance of differences between means was determined by Tukey’s honestly significant difference (HSD) test. Means not sharing the same letter are significantly different (*P* < 0.05). **(B)** *CYP71A12* and *CYP71A13* are dispensable for preinvasive resistance against *Ch*. Aliquots of 2 μL of a conidial suspension of *Ch* (5 × 10^5^ conidia/mL) were drop-inoculated on cotyledons of the tested plants. At 60 hpi, invasive hyphae were observed using light microscopy. In each experiment, at least 50 appressoria were investigated to determine whether they had developed invasive hypha. The means and SDs were calculated from three independent experiments. Statistical analysis using two-tailed Student’s *t* tests showed no significant differences between the genotypes.

### The *bak1–5* mutation reduces the postinvasive resistance of Arabidopsis to *Ab*, independently of pathogen-triggered ICA and camalexin biosynthesis

Next, we investigated the molecular mechanisms of how Arabidopsis recognizes the invasion of necrotrophic pathogens such as *Ab* to activate the biosynthesis of ICAs and camalexin. The candidate mechanism critical for this process is the PRRs-dependent PAMP recognition machinery [36,37]. When Arabidopsis recognizes PAMPs, at least some of the cognate PRRs, including FLS2 and EFR receptor-like kinases (RLKs), form complexes with coreceptors such as BAK1 [27,28,29,30]. Therefore, we decided to assess the possible involvement of BAK1 in the invasion-triggered accumulation of ICAs and camalexin. For this purpose, we used two mutant alleles of *BAK1, bak1–4* and *bak1–5;* of these, *bak1–4* is a *BAK1* null allele [27], and *bak1–5* is a semi-dominant allele with a specific phenotype related to PAMP responsiveness [33]. To precisely compare the results obtained in the assays with *Ab* and with *Ctro* (described below), we used the *pen2 bak1–4* mutant [38] and the newly generated *pen2 bak1–5* mutant. We first performed *Ab* inoculation assay on these plant lines and found that both *pen2 bak1–4* and *pen2 bak1–5* plants have reduced immunity to *Ab*, although the *pen2 bak1– 5* plants were more susceptible than the *pen2 bak1–4* plants (Fig. 5A). Similar results were observed for single *bak1–4* and *bak1–5* as compared with WT plants (Supplementary Fig. S6).

**Figure 5.**
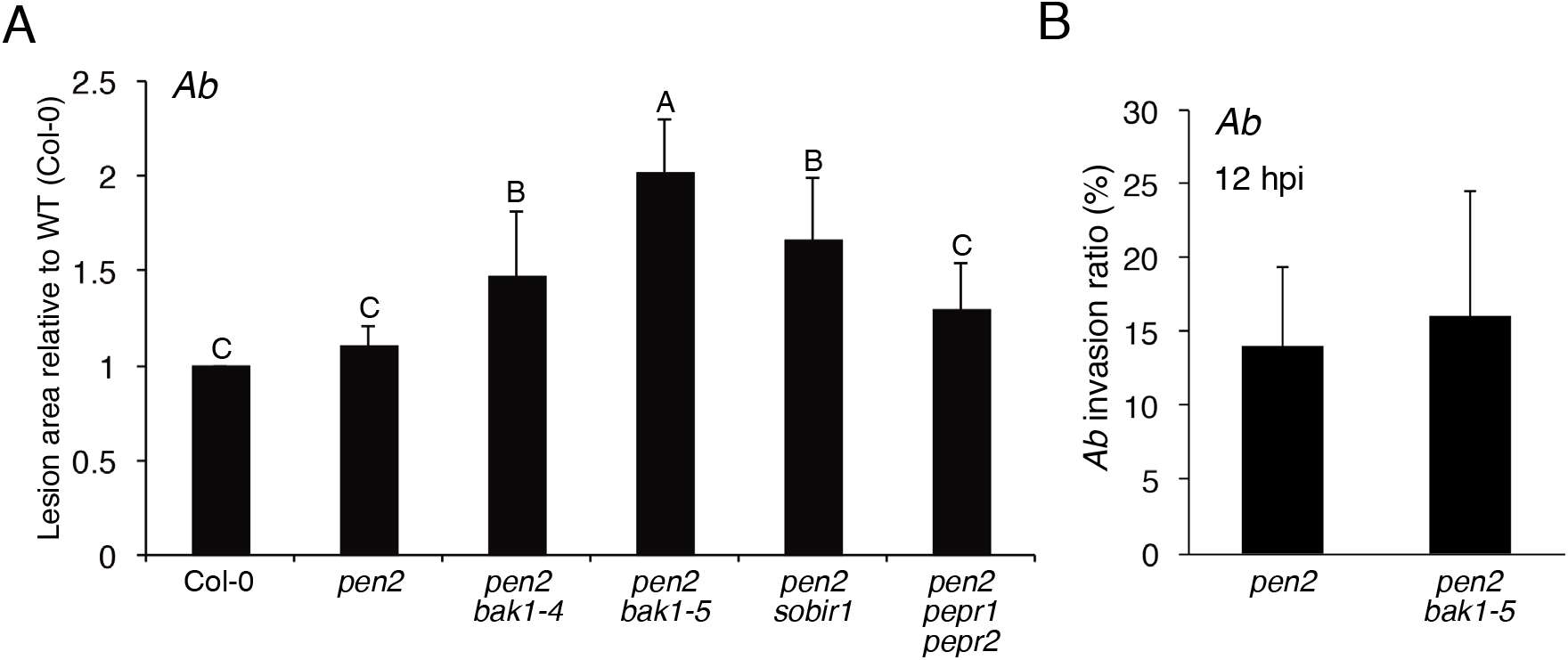
The *bak1* mutations reduced the immunity of Arabidopsis against *Ab*. **(A)** Quantification of lesion development. Conidial suspensions of *Ab* (1 × 10^5^ conidia/mL) were drop-inoculated onto true leaves of 4–5-week-old plants. At 4 dpi, lesion areas were measured and the relative values to Col-0 (WT plants) were calculated. The means and SDs were calculated from three independent experiments. The statistical significance of differences between means was deteπnined by Tukey’s honestly significant difference (HSD) test. Means not sharing the same letter are significantly different (*P* < 0.05). **(B)** The *bak1–5* mutation did not reduce preinvasive resistance against *Ab* in the *pen2* mutant. Aliquots of 5 μL of conidial suspension (1 × 10^5^ conidia/mL) of *Ab* were drop-inoculated onto leaves of 4–5-week-old plants of the tested mutants. At 12 hpi, the inoculated leaves were collected and stained with trypan blue, and then invasive hyphae were observed under light microscopy. At least 50 germinating conidia were counted in each experiment. The means and SDs were calculated from three independent experiments. Statistical analysis using two-tailed Student’s *t* tests showed no significant differences between *pen2* and*pen2 bak1–5* mutants.

We also investigated the possible involvement of SOBIR1 (SUPPRESSOR OF BIR1-1) because this is reported to be essential for triggering defense responses by certain leucine-rich repeat receptor-like proteins (LRR-RLPs) that, together with BAK1, act as immune receptors [39,40,41]. We performed an *Ab* inoculation assay on the *pen2 sobir1* mutant [38] and found that the *pen2 sobir1* mutant has reduced immunity to this fungus (Fig. 5A).

We then evaluated whether the *bak1–5* mutation would reduce preinvasive resistance to *Ab*. To assess this, we compared the invasion behavior of *Ab* in the *pen2* mutant with that in the *pen2 bak1–5* mutant. The invasion ratio in the *pen2 bak1–5* mutant was similar to that in the *pen2* mutant, suggesting that the *bak1–5* mutation does not have detectable impact on preinvasive resistance to *Ab* (Fig. 5B). These results indicate that the *bak1–5* mutation reduces postinvasive resistance to *Ab*, i.e., PRR systems likely function in the recognition of *Ab* invasion. PEPR1 and PEPR2 are LRR-RLKs that recognize the endogenous plant elicitor peptides called Peps [42]. Peps such as AtPep1 are classified as danger-associated molecular patterns (DAMPs) [43]. To assess whether the plant recognition of *Ab* invasion might be related to the possible generation of DAMPs via pathogen invasion, we generated a *pen2 pepr1 pepr2* mutant and performed *Ab* inoculation on this triple mutant; however, we did not find reduced immunity in this triple mutant (Fig. 5A).

Because we found that the *bak1–5* mutation reduced postinvasive resistance to *Ab*, we next investigated whether *bak1–5* would have negative effects on the *Ab* invasion-triggered activation of camalexin and ICA biosynthesis. We checked the gene expression of *PAD3* and *CYP71A12* in the *pen2* and *pen2 bak1–5* mutants after *Ab* inoculation. Surprisingly, both *PAD3* and *CYP71A12* were similarly expressed upon *Ab* invasion in both mutant lines (Fig. 6A). Furthermore, we found that the accumulation of camalexin and ICAs upon *Ab* invasion was not reduced in the *pen2 bak1–5* mutant compared with the *pen2* mutant (Fig. 6B and Supplementary Fig. S7). The accumulation level of ICAs was even higher in *pen2 bak1–5* than *pen2* at 48 hpi, which might be due to enhanced infection in *pen2 bak1–5* (Fig. 6B and Supplementary Fig. S7). Collectively, these results indicate that the *bak1–5* mutation does not reduce the *Ab*-triggered activation of camalexin and ICA biosynthesis, although this mutation reduces postinvasive resistance to *Ab*.

**Figure 6.**
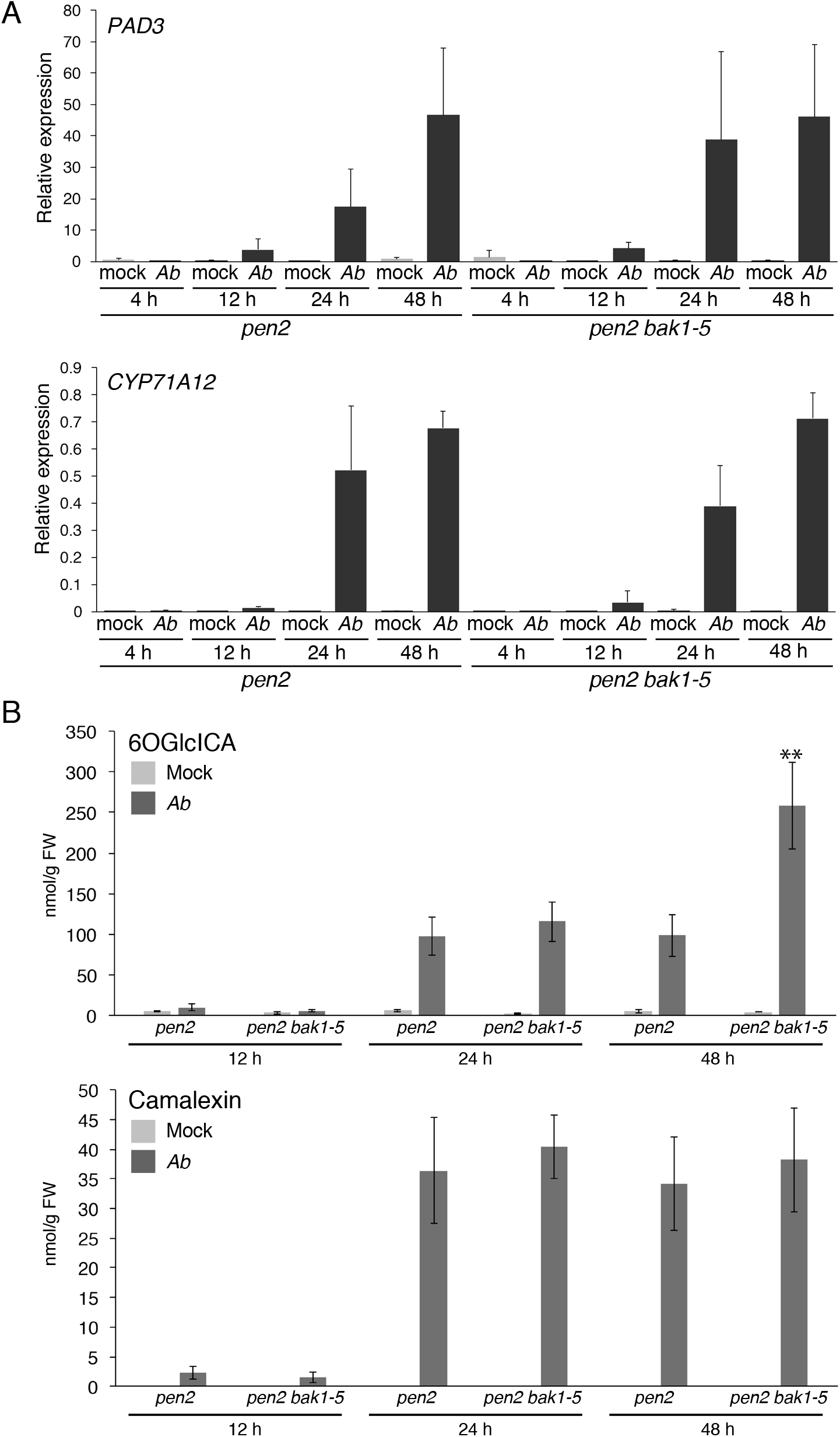
The *bak1–5* mutation did not reduce the *Ab*-invasion triggered accumulation of ICAs and camalexin. **(A)** The *Ab*-invasion triggered activation of *CYP71A12 and PAD3* expression in the *pen2 bak1–5* mutant. Conidial suspensions (5 × 10^5^ conidia/mL) of *Ab* were spray-inoculated onto 4–5-week-old plants. The samples were collected at 4, 12, 24, and 48 hpi. Each gene transcript was quantified by RT–qPCR using the gene-specific primers listed in Supplementary Table S2. Values were normalized to the expression level of *UBC21*. The statistical comparison between 4?-treated *pen2* and *Ab*-treated *pen2 bak1–5* at same time point samples was conducted using two-tailed Student’s t tests and did not show significant differences. **(B)** The *Ab* invasion-triggered accumulations of 6OGlcICA and camalexin were not canceled by the *bak1–5* mutation. Conidial suspensions (5 × 10^5^ conidia/mL) of *Ab* were spray-inoculated onto the tested mutant lines. The means of metabolites (nmol/g FW) and SDs from four biological replicates are shown in the graph. The statistical comparison between *pen2* and *pen2 bak1–5* at same timepoint was conducted using two-tailed Student’s *t* tests (\*\**P* < 0.01).

CHITIN ELICITOR RECEPTOR KINASE 1 (CERK1) is essential for the recognition of fungal chitin [44,45] and triggers immune responses in a BAK1-independent manner [46,47]. Therefore, we assessed whether the activation of Trp-metabolism upon *Ab* infection depends on CERK1-mediated recognition of chitin derived from *Ab*. As a result, we found that the expression of *CYP71A12* and *PAD3* in the *cerk1* mutant upon *Ab* inoculation was comparable to that in WT (Supplementary Fig. S8). These findings suggest that (i) the *bak1–5* mutation impairs or reduces antifungal pathways distinct from Trp metabolism, and that (ii) camalexin and ICA biosynthesis upon *Ab* invasion depend on an unknown pathogen recognition mechanism that is not dependent on BAK1.

### The *bak1–5* mutation reduces the *Ab* invasion-triggered expression of defense-related genes, including *GLIP1*

We further investigated the *bak1–5* sensitive pathways for postinvasive resistance against *Ab*. We performed comparative expression profiling experiments in *pen2* and *pen2 bak1–5* plants following *Ab* invasion using microarray analysis. *Ab* was inoculated to each plant, and RNA isolated from inoculated leaves at 24 hpi was subjected to microarray analysis. We focused on differentially expressed genes associated with the immune response based on Gene Ontology (GO) data (GO term 0006955: http://www.informatics.jax.org). As a result, we found that 14 genes had greater than a 2.5-fold change in expression. Interestingly, 11 were down-regulated, and three were up-regulated (Supplementary Table S1), implying that, compared with the *pen2* plants, the *pen2 bak1–5* plants exhibited a trend for reduced immune responses upon *Ab* invasion. Among the 11 down-regulated genes, four were shown previously to be involved in the Arabidopsis immune system using functional analyses such as the analysis of corresponding knockout mutants, including *AED1* (*APOPLASTIC, EDS1-DEPENDENT 1*) [48], *BGL2/PR2* [49], *GLIP1* [50,51], and *RLP23 (RECEPTOR-LIKE PROTEIN 23*) [40]. Notably, the *glip1* plants were reported to be more susceptible to *Ab* than the WT plant [50]. Subsequently, we performed reverse transcription quantitative polymerase chain reaction (RT–qPCR) analyses to investigate the expression levels of these four genes (*AED1*, *BGL2*/*PR2*, *GLIP1*, and *RLP23*) at 4, 12, 24, and 48 hpi in *pen2* and *pen2 bak1–5* plants inoculated with *Ab* (Fig. 7). As a control, we also investigated gene expression in the mock-treated plants. We confirmed that these four genes exhibited lower expression in the *pen2 bak1–5* than in the *pen2* mutant at 24 hpi, consistent with the array data. Furthermore, the RT–qPCR analysis revealed that the expression of *AED1*, *BGL2*/*PR2*, *GLIP1*, and *RLP23* was lower at 4 hpi than at 12, 24, and 48 hpi (Fig. 7). Together with our finding that *Ab* did not invade at 4 hpi, but started to invade at 12 hpi (Fig. 2B), our results indicate that these genes are induced upon *Ab* invasion to function in postinvasive resistance. Such induced expression was continuously suppressed in the *pen2 bak1–5* plants (Fig. 7), further suggesting that this invasion-triggered expression depends on putative PRRs with function impaired in the *bak1–5* mutant. It is also noteworthy that the invasion-triggered expressions of *AED1*, *BGL2/PR2, GLIP1*, and *RLP23* were time-dependent, i.e., induced expression started to be down-regulated at 12 hpi (*RLP23*) or 24 hpi (*AED1*, *BGL2*/*PR2*, *GLIP1*) (Fig. 7), which is in contrast to the invasion-triggered expressions of *PAD3* and *CYP71A12*, which exhibited sustained elevations for up to 48 hpi (Figs. 2 and 6).

**Figure 7.**
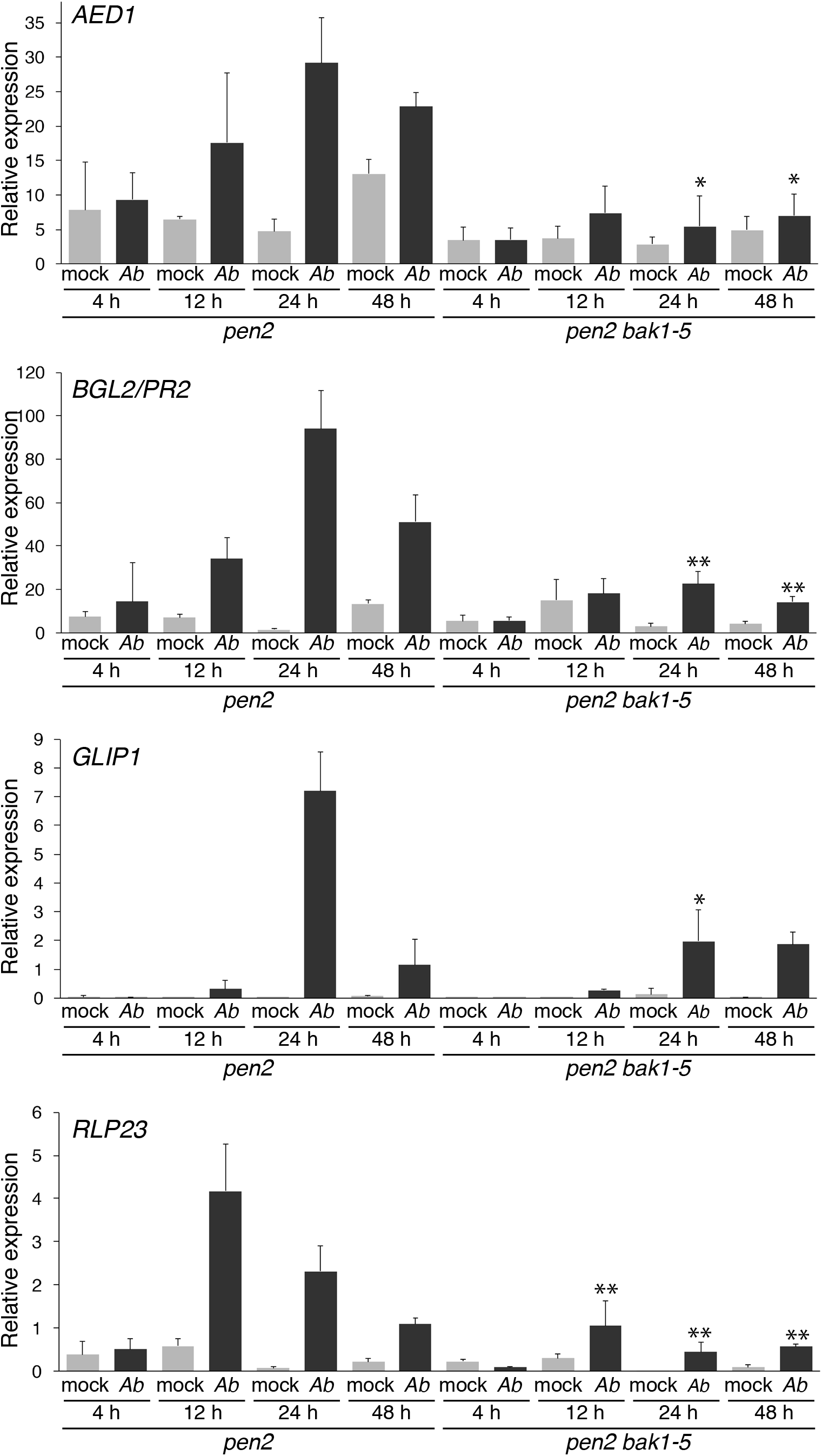
The *Ab* invasion-triggered expression levels of *AED1, BGL2, GLIP1*, and *RLP23* were reduced by the *bak1–5* mutation. Conidial suspensions (5 × 10^5^ conidia/mL) of *Ab* were spray-inoculated onto 4–5-week-old*pen2* and *pen2 bak1–5* plants, and then kept at 100% humidity. The samples were collected at 4, 12, 24, and 48 hpi. Each gene transcript was quantified by RT–qPCR using the gene-specific primers listed in Supplementary Table S2. Values were noπnalized to the expression level of *UBC2l*. The means and SDs were calculated from three independent experiments. The statistical analysis was conducted by a two-tailed *t* test. The expression levels of each gene between *pen2* and*pen2 bak1–5* were compared at the same time points and treatment (\**P* < 0.05; \*\**P* < 0.01).

### Conserved machinery in the immunity of Arabidopsis toward invasion of multiple fungal pathogens with distinct infection modes

Up to here, we have revealed that postinvasive resistance toward *Ab* involves (i) a Trp-related secondary metabolism including the biosynthesis of both camalexin and ICAs, and (ii) a Trp-unrelated defense pathway that is sensitive to the *bak1–5* mutation. Arabidopsis exhibits robust nonhost resistance against the nonadapted hemibiotroph *Ctro*, and full postinvasive resistance against *Ctro* requires camalexin and ICAs synthesis, which requires induced expression of *CYP71A12, CYP71A13* and *PAD3* [23]. Therefore, we tested whether the postinvasive resistance against *Ctro* would also be affected by the *bak1–5* mutation. We reported previously that the *bak1–5* mutation reduces preinvasive resistance to *Ctro* [52]. Here, a trypan blue staining viability assay revealed that the *bak1–5* mutation also reduces postinvasive resistance to this nonadapted pathogen (Fig. 8A). We can exclude the possibility that reduced preinvasive resistance in the *pen2 bak1–5* mutant results in decreased postinvasive resistance to *Ctro*, because the *pen2 edr1* plants exhibited a more severe reduction in preinvasive resistance, but still had no detectable reduction in postinvasive resistance to *Ctro* [16]. Thus, *bak1–5* reduced postinvasive resistance toward both *Ab* and *Ctro*.

**Figure 8.**
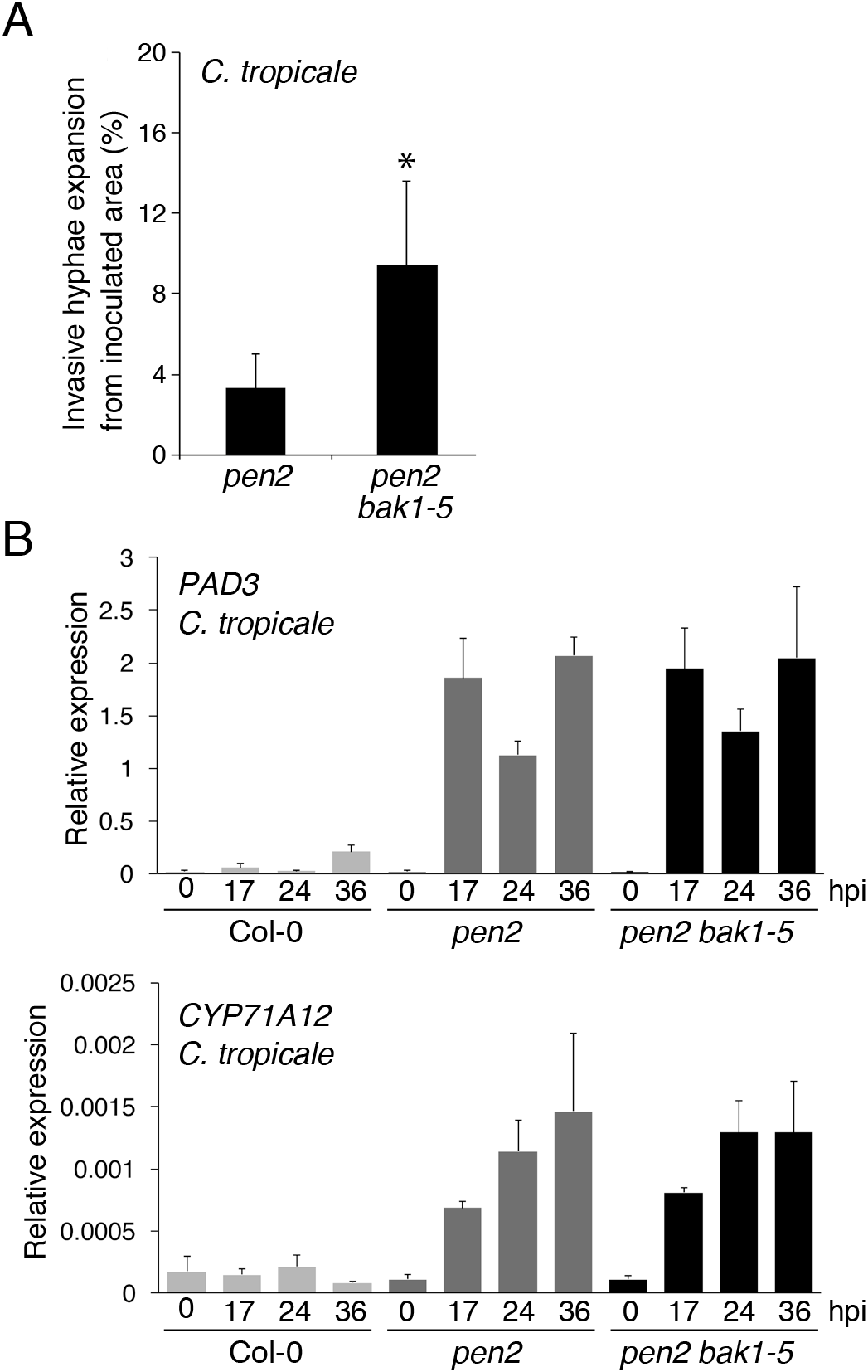
The *bak1–5* mutation reduced postinvasive resistance against the nonadapted hemibiotrophic pathogen *Colletotrichum tropicale* (*Ctro*), but did not affect the *Ctro*-invasion-triggered expression levels of *CYP71A12* or *PAD3*. **(A)** The *bak1–5* mutation reduced postinvasive resistance against *Ctro*. Aliquots of 5 μL of conidial suspension (2.5 × 10^5^ conidia/mL) of *Ctro* were drop-inoculated onto the tested plants. At 4 dpi, the inoculated leaves were collected and stained with Trypan blue. The expansion of invasive hyphae from the inoculated area was investigated using light microscopy, and the percentages of expanded invasive hyphae were determined. The means and SDs were calculated from three independent experiments. The statistical analysis was conducted using two-tailed Student’s *t* tests (\*\**P* < 0.01). **(B)** The *Ctro*-invasion-triggcrcd expression of *PAD3* and *CYP71A12* was not canceled by the *bak1–5* mutation. Conidial suspensions (5 × 10^5^ conidia/mL) of *Ctro* were spray-inoculated onto the tested plants. Leaf samples were collected at 17, 24, and 36 hpi. Each gene transcript was quantified by RT– qPCR using the gene-specific primers listed in Supplementary Table S2. Values were normalized to the expression level of *UBC21*. The means and SDs were calculated from three independent experiments. Statistical analysis using two-tailed Student’s *t* tests showed no significant differences between the *pen2* and*pen2 bak1–5* mutants at the same time points.

We further investigated whether the *bak1–5* mutation would have negative effects on the expression of *CYP71A12* and *PAD3* triggered by *Ctro* invasion. We observed that the expressions of *CYP71A12* and *PAD3* were induced in *pen2* but not in WT plants, indicating that their expressions were triggered by *Ctro* invasion (Fig. 8B); notably, we found that the *bak1–5* mutation had no clear effect on this (Fig. 8B). Thus, postinvasive resistance against *Ctro* also requires (i) a *bak1–5*-insensitive Trp-metabolism including synthesis of camalexin and ICAs, and (ii) a *bak1–5*-sensitive defense response uncoupled from Trp-metabolism, which is quite similar to postinvasive resistance against *Ab* (Fig. 9). Collectively, these results represent unexpectedly strong conservation of machineries for postinvasive resistance against two distantly related fungal pathogens having distinct infection strategies, i.e., a brassica-adapted necrotrophic fungus and a nonadapted hemibiotrophic fungus.

**Figure 9.**
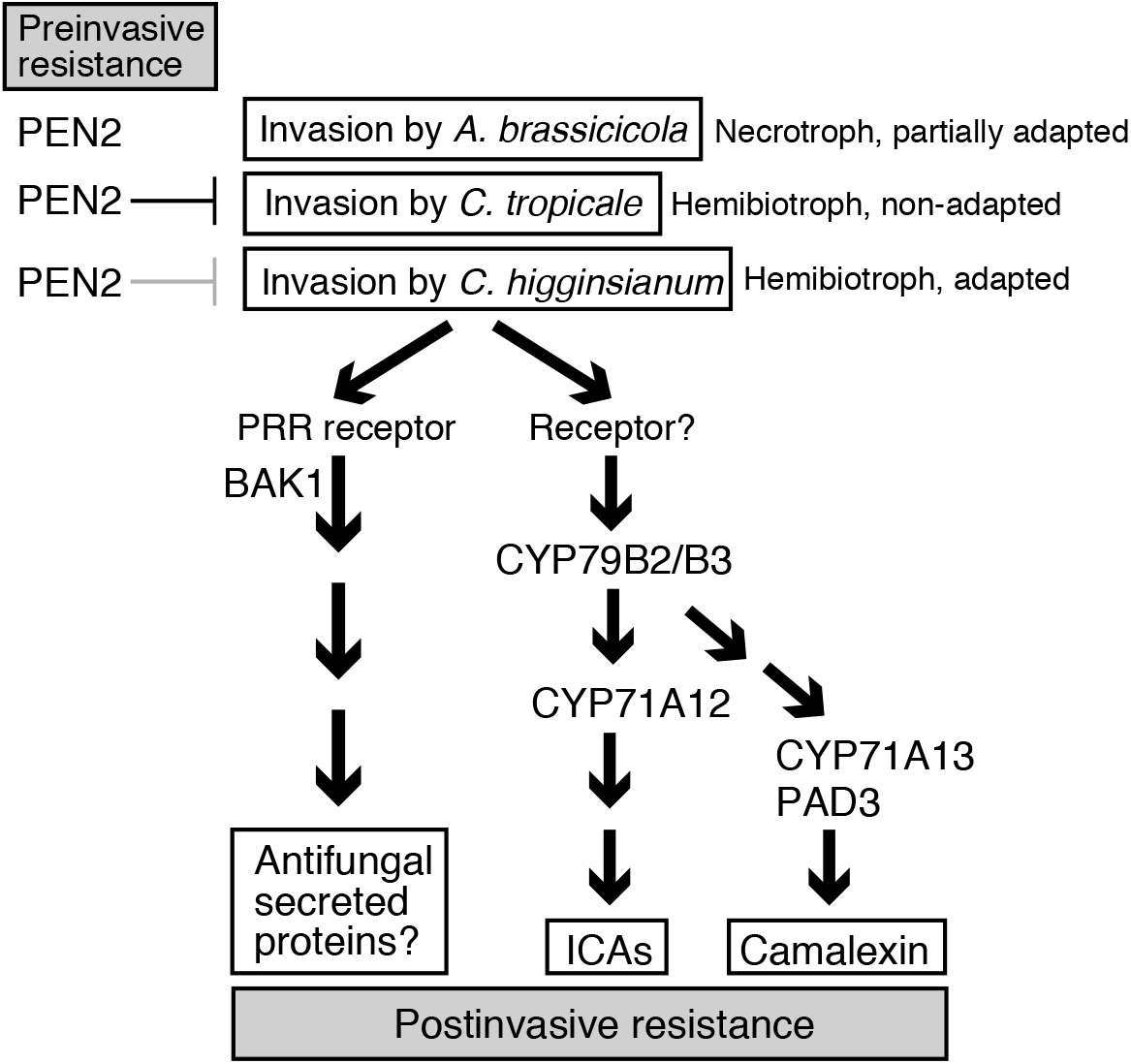
Summarized model for Arabidopsis postinvasive resistance against fungal pathogens. The CYP7lAl2-dependent production of ICAs is required for postinvasive resistance against *A. brassicicola, C. tropicale*, and *C. higginsianum* with different infection modes. The CYP71A13- and PAD3-dependent production of camalexin is also required for postinvasive resistance against the three pathogens. The *bakl* mutations (especially *bak1–5*) reduced the postinvasive resistance, however, invasion-triggered activation of these Trp-related pathways is not canceled by *bak1–5*. The *bak1–5* sensitive pathway control the expression of antifungal protein genes (ex. *GLIP?*). In contrast to conserved machineries for postinvasive resistance against different pathogens, factors required for preinvasive resistance are diverse among pathogens. Different contribution of PEN2 to preinvasive resistance against the three pathogens is shown here as an example.

## DISCUSSION

We recently reported that CYP71A12 is indispensable for postinvasive resistance to the hemibiotrophic pathogen *Ctro* that is not adapted to Arabidopsis [23]. However, it remains obscure whether CYP71A12 is also involved in this invasion-triggered resistance of Arabidopsis against other fungal pathogens.

Here we found that the absence of functional CYP71A12 enhanced lesion development during colonization of Arabidopsis leaves with the necrotrophic pathogen *Ab*, indicating that this enzyme is required for the immune response of Arabidopsis against this fungus. *CYP71A12* was not induced at 4 hpi with *Ab* when the pathogen had not yet invaded, but it started to be induced upon *Ab* invasion. We also found that *CYP71A12* was dispensable for preinvasive resistance against *Ab*. Collectively, these results demonstrate that CYP71A12 is required for postinvasive resistance against *Ab* as well as *Ctro*. Metabolic analyses showed that the enhanced accumulation of ICAs triggered by *Ab* invasion was reduced in the absence of functional CYP71A12. Thus, this enzyme is involved in the accumulation of ICAs following *Ab* invasion (Fig. 3B). Therefore, we consider that the CYP71A12-dependent synthesis of ICA and/or its derivatives upon *Ab* invasion is critical for postinvasive resistance to this fungus.

We also showed that CYP71A13 contributes to the postinvasive resistance against *Ab* via synthesis of camalexin, but not of ICAs. The result suggests the importance of camalexin for this second layer of defense, which is further supported by our phenotypic analyses of mutants defective in *PAD3* (Fig. 1 and Supplementary Fig. S1). Remarkably, we also found that CYP71A12, CYP71A13, and PAD3 are involved in the postinvasive resistance of Arabidopsis against *Ch*, an adapted hemibiotrophic fungus. These results strongly suggest the broad importance of camalexin and ICAs for postinvasive resistance against fungal pathogens with distinct infection strategies.

By contrast, PEN2 is involved in preinvasive resistance against *Ctro* and *Ch* [11,14], but dispensable for preinvasive resistance against *Ab* (Fig. 2D). Also, PEN2 is dispensable for nonhost resistance against non-adapted isolates of *P. cucumerina*, but is involved in basal resistance against an adapted isolate of this necrotrophic fungus, however, it remains to be elucidated whether this involvement results from the PEN2 function in preinvasive resistance [21]. Furthermore, PEN1 is known to be required for preinvasive resistance against nonadapted powdery mildews, but dispensable for the *Colletotrichum* fungi and nonadapted *Alternaria alternata* [13,53]. Thus, the molecular components that underlie preinvasive resistance vary between fungal pathogens, probably because pathogens have evolved various strategies for plant entry (Fig. 9). By contrast, once these pathogens enter Arabidopsis, they commonly develop invasive structures inside the plants; thus, the hosts might deploy an universal defense systems to terminate further fungal growth (Fig. 9).

We found that the *cyp79B2 cyp79B3* mutant is more susceptible to *Ab* than is the *pen2 cyp71A12 cyp71A13* mutant (Fig. 1 and Supplementary Fig. S2). Interestingly, this phenomenon is also observed in the infection by *Ch* (Fig. 4A), *Ctro*, and *P. cucumerina* [23]. Metabolite analyses of plants upon *Ab* invasion revealed that camalexin accumulation in the *pen2 cyp71A12 cyp71A13* plants was almost the same as that in the *cyp79B2 cyp79B3* plants. Importantly, the *pen2 cyp71A1 cyp71A13* plants reduced their accumulation of ICAs, but still produced them to some degree, whereas the *cyp79B2 cyp79B3* plants were entirely defective in this regard. Consistent with this finding, several independent metabolic branches are supposed to contribute to endogenous ICAs levels, including contribution of AAO1 (Arabidopsis aldehyde oxidase 1) and CYP71B6 [23,54,55].

We also compared the phenotype of *pen2 cyp71A12 cyp71A13* mutants with that of *myb34 myb51 myb122 cyp71A12 cyp71A13* following *Ab* invasion and found no detectable differences in either mutants in terms of postinvasive resistance against *Ab*, indicating that the difference between *pen2 cyp71A12 cyp71A13* and *cyp79B2 cyp79B3* mutants is not caused by PEN2-unrelated IG metabolism products (Supplementary Fig. S5). Collectively, these findings suggest that the lower susceptibility in the former mutants is caused by residual ICAs, i.e., ICAs contribute to postinvasive resistance in a dose-dependent manner. This supports the idea that ICAs or their derivatives work as antifungal compounds as opposed to functioning as signaling molecules for plant immune responses. However, we cannot exclude a possibility that accumulation of so far not-reported IAOx-derivatives whose biosynthesis is not dependent on CYP71A12 and CYP71A13 contribute to the difference observed in susceptibility of *pen2 cyp71A12 cyp71A13* and *cyp79B2 cyp79B3* plants.

We also investigated how Arabidopsis recognizes the invasion of fungal pathogens and then mounts its postinvasive resistance. We found that the two *bak1* mutations, especially the *bak1–5* mutation, reduce postinvasive resistance against *Ab* (Fig. 5A and Supplementary Fig. S6). The finding that the negative effects of *bak1-5* on the postinvasive resistance towards *Ab* was higher than that of *bak1-4* is likely consistent with the previous works [33].

Surprisingly, we found that the *bak1–5* mutation did not hamper the invasion-triggered expressions of *PAD3* and *CYP71A12* and the subsequent accumulation of ICAs and camalexin (Fig. 6). Thus, we postulate existence of another defense mechanism that is affected by the *bak1–5* mutation and required for postinvasive resistance against *Ab*. Our further analyses revealed that this pathway activates the expression of distinct defense-related genes, including *AED1*, *BGL2/PR2*, *GLIP1*, and *RLP23* (Supplementary Table S1 and Fig. 7). Notably, expressions of these genes were induced following *Ab* invasion, and these were canceled in *bak1–5* mutants (Fig. 7). Therefore, we suggest that Arabidopsis deploys a common PRR system to sense the invasion of multiple fungal pathogens and subsequently activate antifungal defense pathways that are uncoupled from Trp-metabolism (Fig. 9). We hypothesized that sensing pathogen invasion involves the recognition of DAMPs. It is known that Arabidopsis perceives endogenous Pep peptides by two redundant PRRs: PEPR1 and PEPR2 [41]. Moreover, the PEPR1/PEPR2 pathway is impaired by the *bak1–5* mutation [56]. However, we found that immunity to *Ab* was not significantly reduced in the *pepr1 pepr2* mutant (Fig. 5A). It will therefore be important to identify the corresponding PRR in the *bak1–5*-sensitive pathway for postinvasive resistance.

Notably, it has been reported that the *glip1* mutant plants exhibit enhanced susceptibility to *Ab* [50]. The recombinant GLIP1 protein exhibits antimicrobial activity that disrupts the *Ab* spores and hyphae [50] and triggers systemic acquired resistance (SAR) against bacterial pathogens (e.g., *Erwinia carotovora* and *Pseudomonas syringae*) as well as *Ab* [51]. Thus, the enhanced susceptibility of Arabidopsis to *Ab* in the presence of *bak1–5* might be partially caused by the reduced expression of *GLIP1*.

It remains unclear how this plant recognizes pathogen invasion and then activates Trp-related metabolite accumulation as key immune responses in postinvasive resistance. Our data suggest that the corresponding pathway is not sensitive to the *bak1– 5* mutation. We also found that the *Ab*-triggered activation of Trp-related metabolism does not depend on CERK1 that is autonomous from BAK1 (Supplementary Fig. S8). Because *PAD3* and *CYP71A12* were commonly induced by the invasion of diverse fungal pathogens such as *Ab* and *Ctro*, we suggest that Arabidopsis probably recognizes the cell damage that is commonly caused by pathogen invasion and then activates Trp-related metabolism. Further studies are needed to explore a recognition mechanism of pathogen invasion that activates this secondary metabolic pathways for antifungal defense.

## MATERIALS AND METHODS

### Fungal materials

*C. tropicale (Ctro*) (formerly *Colletotrichum gleosporioides* S9275) was provided by Shigenobu Yoshida (National Institute for Agro-Environmental Sciences, Japan); *C. higginsianum* (*Ch*) isolate MAFF305635 was obtained from the Ministry of Agriculture, Forestry and Fisheries (MAFF) Genebank, Japan; and *A. brassicicola* (*Ab*) strain Ryo-1 was provided by Akira Tohyama. Cultures of *Ch* and *Ab* were maintained on 3.9% (w/v) potato dextrose agar medium (PDA; Nissui Pharmaceutical Co., Ltd., Tokyo, Japan) at 24 °C in the dark. *Ctro* was cultured on 2.5% (w/v) PDA (Difco, Detroit, MI, USA) at 24 °C under a cycle of 16 h black light (FS20S/BLB 20W; Toshiba, Tokyo, Japan) illumination and 8 h dark.

### Arabidopsis lines and growth conditions

The *A. thaliana* accession Col-0 was used as the WT plant. The mutants *pen2-1*, *pen2-2* [3], *pad3-1* [57], *cyp71A12, cyp71A12/cyp71A13* [60], *cyp82C2* [20], *bak1–4* [27], *bak1–5* [33], *cyp79B2 cyp79B3* [17], *pen2 pad3* [5,] *pen2 cyp82C2* [23], *pen2 cyp71A12* [23], *pepr1 pepr2* [42], *pen2 cyp71A12 cyp71A13* [23], *pen2 sobir1* [38], *myb34 myb51 myb122* [23], *cyp71A12 cyp71A13 myb34 myb51 myb122* [23] and *cerk1-2* [44] were used in this study. Arabidopsis seeds were sown on rockwool (Grodan; http://www.grodan.com) and kept at 4 °C in the dark for 2 days, and later grown at 25 °C with a cycle of 16 h light and 8 h dark in Hoagland medium.

### Pathogen inoculation, lesion development analysis and trypan blue viability staining assay

For spray inoculation assays of *Ab*, *Ch*, and *Ctro*, 5 × 10^5^ conidia/mL of conidial suspension was spray-inoculated on 4–5-week-old plants. For drop-inoculation, 5 μL of conidial suspensions of *Ab*, *Ch* (1 × 10^5^ conidia/mL) or *Ctro* (2.5 × 10^5^ conidia/mL) were placed onto each leaf. Conidia of *Ctro* were inoculated with 0.1% (w/v) glucose. The inoculated plants were kept at 25 °C with a cycle of 16 h light and 8 h dark and maintained at 100% relative humidity. For analysis of lesion development following the inoculation of *Ab* or *Ch*, four drops of 5 μL conidial suspension of each pathogen were drop-inoculated on each leaf, and 24–50 lesions were evaluated in each experiment. The developed lesions were quantified using ImageJ image analysis software (http://imagej.net) and relative values to WT (Col-0) plants were calculated. To measure lesion areas, yellowish areas were included as lesions. Trypan blue staining was conducted according to [59]. For the trypan blue assay, at least 50 lesions were investigated in each experiment. The *Ab* invasion ratio (%) was calculated by using the following formula: *Ab* invasion ratio (%) = (number of germinating conidia that developed invasive hyphae / number of germinating conidia) × 100. For the *Ch* invasion assay, 2 μL aliquots of 5 × 10^5^ conidia/mL of the *Ch* conidial suspension were drop-inoculated onto Arabidopsis cotyledons (14 days old). At 60 hpi, invasive hyphae were investigated using light microscopy. The invasion ratio of *Ch* was calculated as described previously [52].

### Generation of mutant plants

The generation of *pen2 bak1–4*, *pen2 bak1–5* and *pen2 pepr1 pepr2* lines used in this study were generated by crossing the *pen2–1* mutant with *bak1–4*, *bak1–5*, or *pepr1 pepr2* plants. Each genotype was checked with the corresponding specific primers for the derived cleaved-amplified polymorphic sequence (dCAPS) markers using dCAPS Finder 2.0 (http://helix.wustl.edu/dcaps/dcaps.html), and the PCR products (WT or mutant types) were cleaved with appropriate restriction enzymes (Supplementary Table S2).

### RT–qPCR analysis

Seven Arabidopsis leaves inoculated with *Ab* or *Ch* (5 × 10^5^ conidia/mL) were collected from each of seven different plants of either WT Col-0 or mutant plants at corresponding time points. Total RNA was extracted using PureLink (TRIzol plus RNA purification kits, Life Technologies/Thermo Fisher Scientific, Waltham, MA, USA) and treated with DNase (RQ1 RNase-free DNase; Promega, Madison, WI, USA; http://www.promega.com) to remove DNA contamination. Takara Prime Script™ RT kits (Takara Bio Inc., Shiga, Japan; http://www.takara-bio.com) was used for the cDNA synthesis. Takara TB Green™ Premix Ex Taq™ I was used for RT–qPCR, performed using the primers listed in Supplementary Table S2. Arabidopsis *UBC21* (At5g25760) was used as an internal control for normalizing the level of cDNA [60]. RT–qPCR analysis was performed using a Thermal Cycler Dice Real Time System TP800 (Takara). The expression levels of genes of interest were normalized relative to those of *UBC21*. Relative expression (log2) was calculated by subtracting Ct values of genes of interest from those of *UBC21*. Fold change was based on values of relative expression (log2), which was calculated by two to the power of relative expression (log2). For the statistical analysis, relative expression values (in log2 scale) were used, whereas the graphs of the RT–qPCR analysis in figures were represented using fold change values (for easy observation).

### Metabolite analysis

Conidial suspensions (5 × 10^5^ conidia/mL) of *Ab* were spray-inoculated onto 4–5-week-old plants and kept at 100% relative humidity. Leaf samples (100–200 mg fresh weight) were collected at corresponding time points and frozen immediately in liquid nitrogen. The plant extracts containing Trp derivatives were extracted using DMSO and metabolite analyses were performed as described [5,23].

### Microarray analysis

*Ab* conidial suspensions (5 × 10^5^ conidia/mL) were spray-inoculated onto 4–5-week-old plants of the *pen2* and *pen2 bak1–5* mutants. For each sample, five leaves were collected at 24 hpi and frozen immediately in liquid nitrogen. In total, eight samples (four biological replicates each of the mock and *Ab*-treated samples) were used for RNA extraction. Total RNA was extracted using Plant RNA Isolation Mini kits (Agilent Technologies., Santa Clara, CA, USA). Aliquots of 200 ng of total RNA were used to prepare Cy3-labeled cRNA using Agilent Low Input Quick Amp labeling kits. The labeled samples were hybridized onto an Agilent Arabidopsis *thalian*a microarray (ver. 4.0; 4 · 44 K format). After hybridization and washing, the arrays were scanned using an Agilent microarray scanner (G2565BA). The images were analyzed using Agilent Feature Extraction software (ver. 10.7.3.1), and further analysis was performed using Agilent GeneSpring GX12.1 software. Signal normalization was based on the expression ratio of *pen2 bak1–5* to *pen2*. Differentially upregulated genes were defined as having a greater than 2.5-fold increase in expression, and differentially downregulated genes were defined as having at least a 0.4-fold decrease in expression. Microarray data have been deposited in the NCBI Gene Expression Omnibus (GEO) database GSE 124921.

## ACKNOWLEDGEMENTS

We thank Yube Yamaguchi (the *pepr1 pepr2*) and ABRC for providing Arabidopsis seeds. We also thank Shigenobu Yoshida (*Ctro*), Akira Tohyama (*Ab*), and the MAFF Genebank (*Ch*) for providing fungal pathogens. We also thank Yoshihiro Inoue for supports on the statistical analyses. This work was supported by Grants-in-Aid for Scientific Research (15H05780, 18H02204, 18H04780, 18K19212) (KAKENHI), by grants from the Project of the NARO Bio-oriented Technology Research Advancement Institution (Research program on development of innovative technology), and by the Asahi Glass Foundation. Work in the PB laboratory was supported by the National Science Centre SONATA BIS grant (UMO-2012/07/E/NZ2/04098).

## AUTHORS CONTRIBUTIONS

Y.T. and P.B. designed this research. A.K., M.P., M.P-B., T.N., H.S., A.I. and H.F. performed the experiments and analyzed the data. A.K., Y.T., P.B., M.K. and K.M. wrote the manuscript and prepared the figures.

## Notes

### Competing Interest Statement

The authors have declared no competing interest.

