## Supplemental Information for "*bak1-5* mutation uncouples tryptophan-dependent and independent postinvasive immune pathways triggered in Arabidopsis by multiple fungal pathogens"

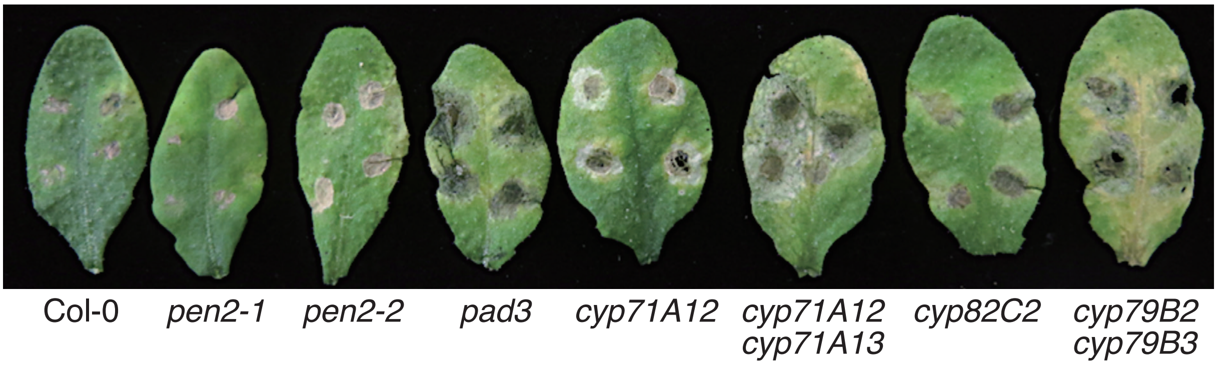


**Supplementary Fig. S1.** Lesion development caused by *Ab* on Arabidopsis mutant plants. Conidial suspensions (1 x $10^5$ conidia/mL) of *Ab* were drop-inoculated onto the tested mutant lines. This photograph was taken at 4 days after the inoculation.

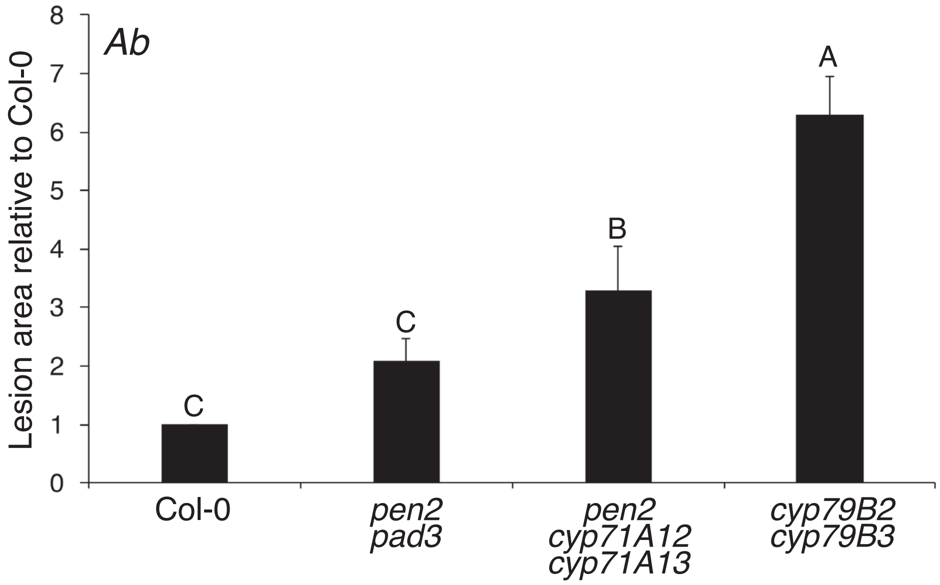


**Supplementary Fig. S2.** The *cyp79B2 cyp79B3* mutant was more susceptible to *Ab* than was the *pen2 cyp71A12 cyp71A13* mutant. Quantification of lesion development in the *Ab*-inoculated leaves of the tested Arabidopsis lines at an early time point (3 dpi). Conidial suspensions of *Ab* ($1 \times 10^5$ conidia/mL) were drop-inoculated onto true leaves of 4–5-week-old plants. At 3 dpi, lesion areas were measured, and then the relative values to those in Col-0 (WT plants) were calculated. The means and SDs were derived from three independent experiments. The statistical significance of differences between means was determined by Tukey's honestly significant difference (HSD) test. Means not sharing the same letter are significantly different ($P < 0.05$).

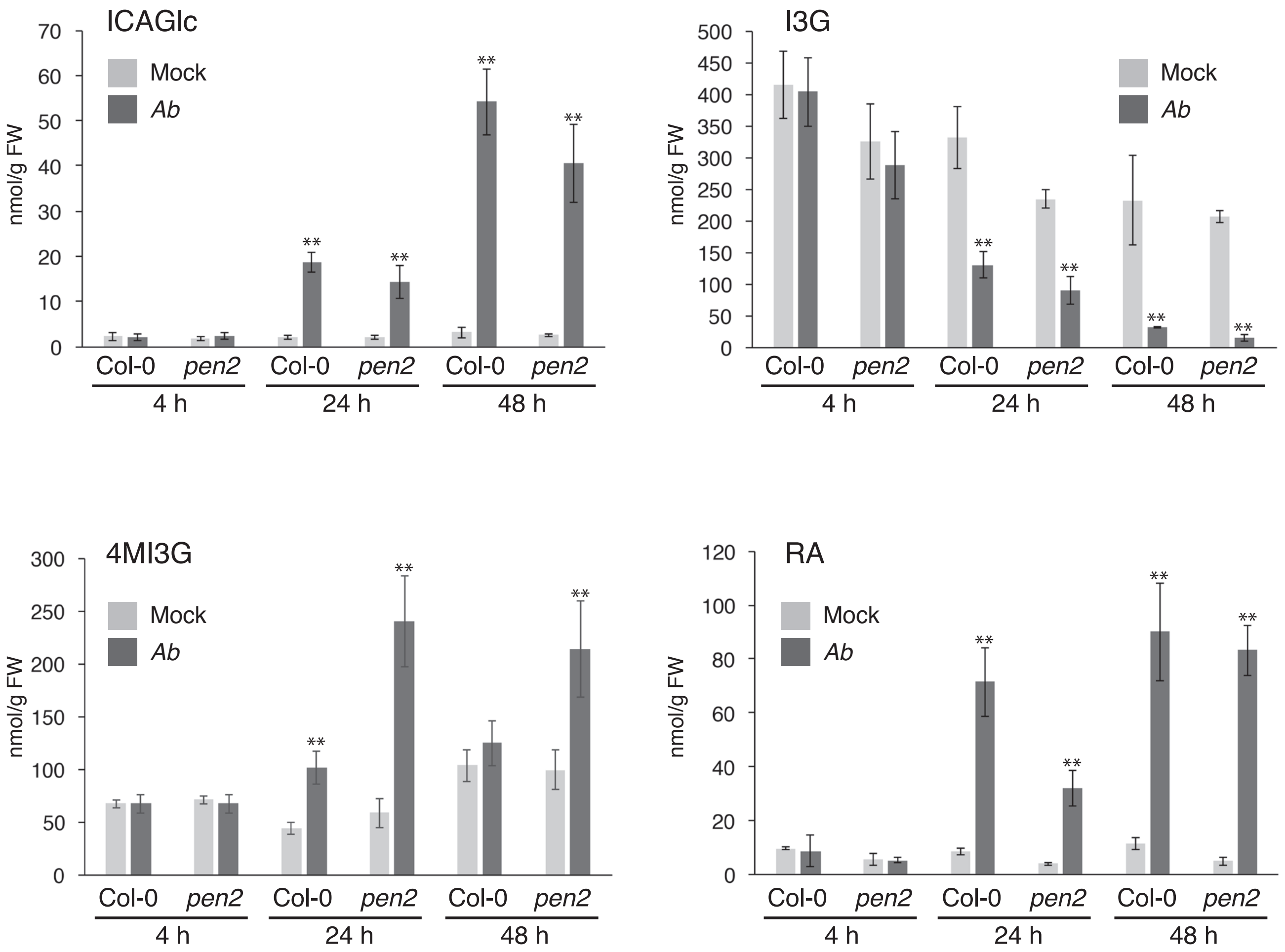


**Supplementary Fig. S3.** The analysis of ICAGlc, I3G, 4MI3G, and RA accumulation in WT (Col-0 plants) and the *pen2* mutant inoculated with *Ab*. Conidial suspensions (5 x $10^5$ conidia/mL) of *Ab* were spray-inoculated onto WT and *pen2* plants. As a control, water was sprayed as a mock treatment. The samples were collected at 4, 24, and 48 hpi. The accumulation of indole-3-carboxylic acid glucose ester (ICAGlc), indole-3-ylmethylglucosinolate (I3G), 4-methoxyindol-3-ylmethylglucosinolate (4MI3G), and raphanusamic acid (RA) were determined. The means of metabolites (nmol/g FW) and SDs from four biological replicates are shown in the graph. The statistical analysis between mock- and *Ab*-treated samples at each time point was conducted using two-tailed Student's *t* tests (**$P < 0.01$).

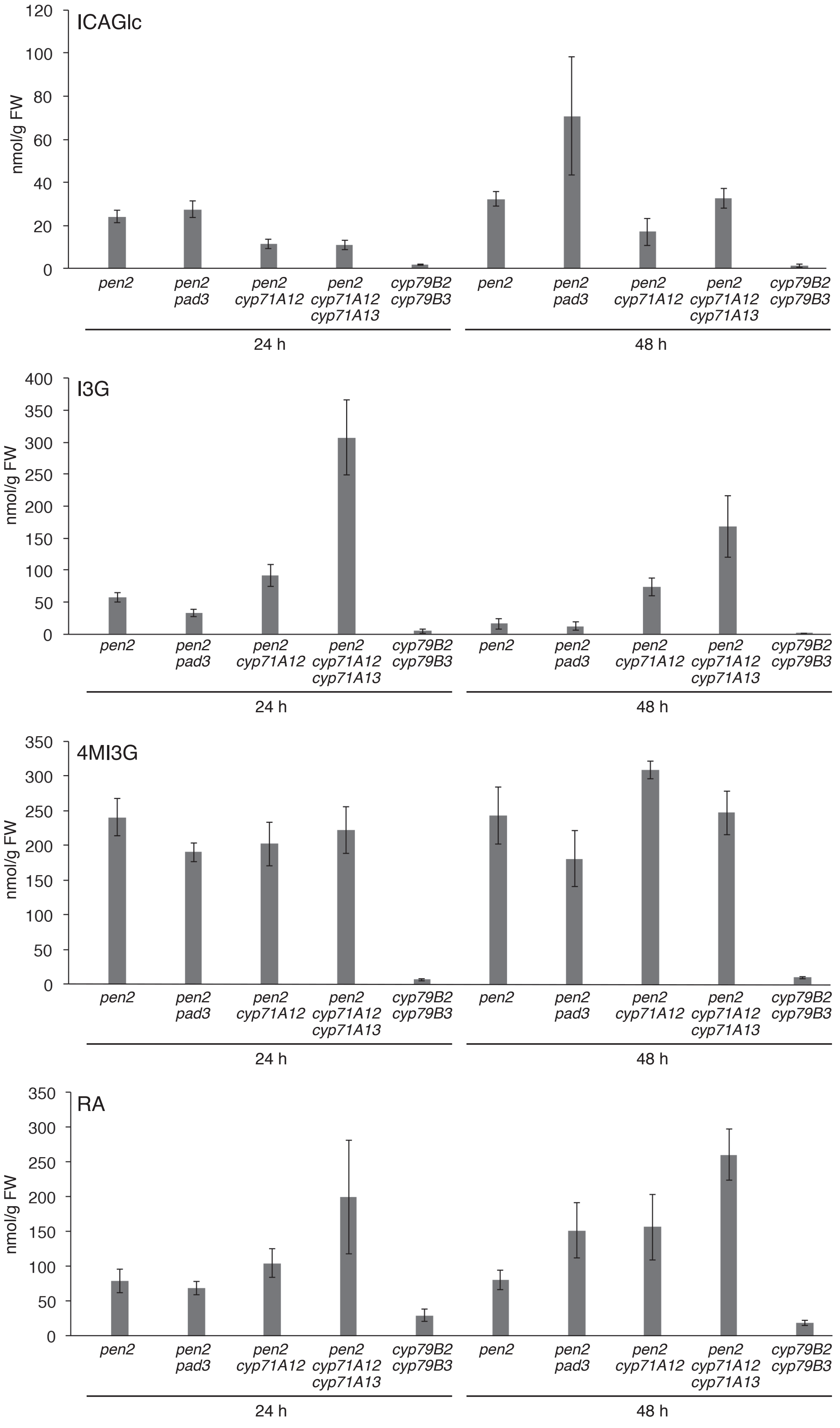


**Supplementary Fig. S4.** The analysis of ICAGlc, I3G, 4MI3G, and RA accumulation in a series of mutants defective in Trp metabolism. Conidial suspensions (5 x $10^5$ conidia/mL) of *Ab* were spray-inoculated onto the tested mutant plants. The samples were collected at 24 and 48 hpi. The accumulations of ICAGlc, I3G, 4MI3G, and RA were determined. The means of metabolites (in nmol/g FW) and SDs from four biological replicates are shown in the graph.

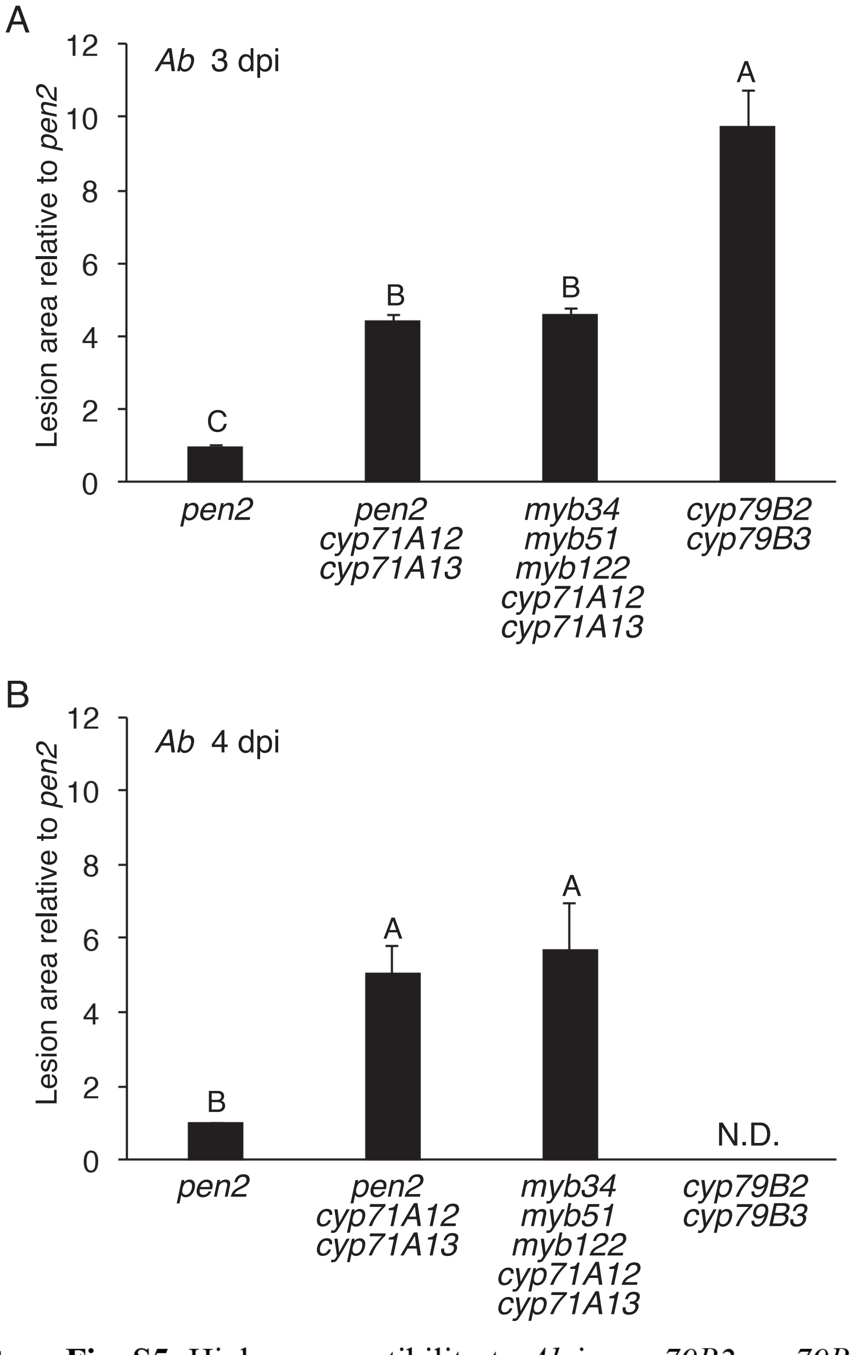


**Supplementary Fig. S5.** Higher susceptibility to *Ab* in *cyp79B2 cyp79B3* than in *pen2 cyp71A12 cyp71A13* mutants is not caused by the accumulation of PEN2-independent indole-glucosinolates. Conidial suspensions of *Ab* ($1 \times 10^5$ conidia/mL) were drop-inoculated onto true leaves of 4–5-week-old plants. At 3 dpi (A) and 4 dpi (B), lesion areas were measured, and relative values to the *pen2* mutant were calculated. The means and SDs were derived from three independent experiments. The statistical significance of differences between means was determined by Tukey's honestly significant difference (HSD) test. Means not sharing the same letter are significantly different ($P < 0.05$). ND; not determined.

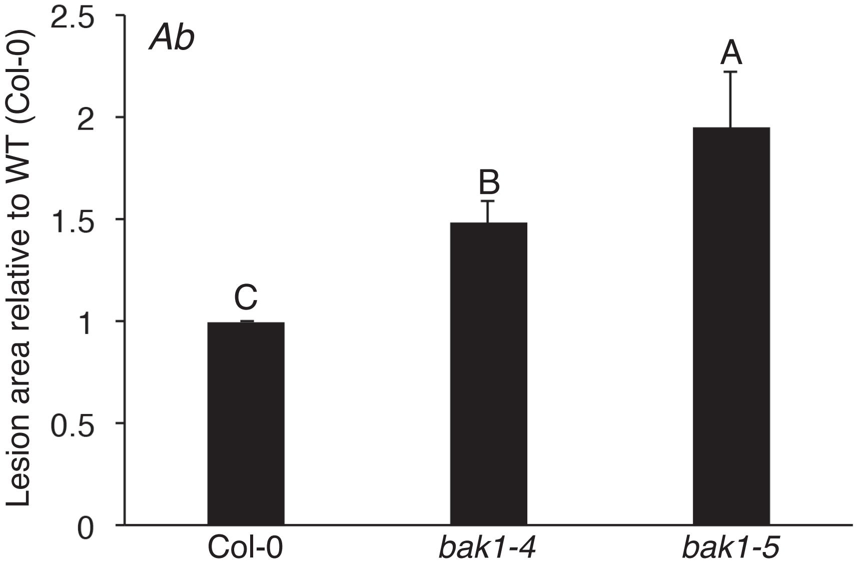


**Supplementary Fig. S6.** The *bak1* mutants showed enhanced susceptibility toward *Ab* compared with the WT plants (Col-0). Conidial suspensions of *Ab* (1 x $10^5$ conidia/mL) were drop-inoculated onto the tested plants. At 4 dpi, lesion development relative to WT plants was determined. The means and SDs were derived from three independent experiments. The statistical significance between the means was determined by Tukey's honestly significant difference (HSD) test. Means not sharing the same letters are significantly different ($P < 0.05$).

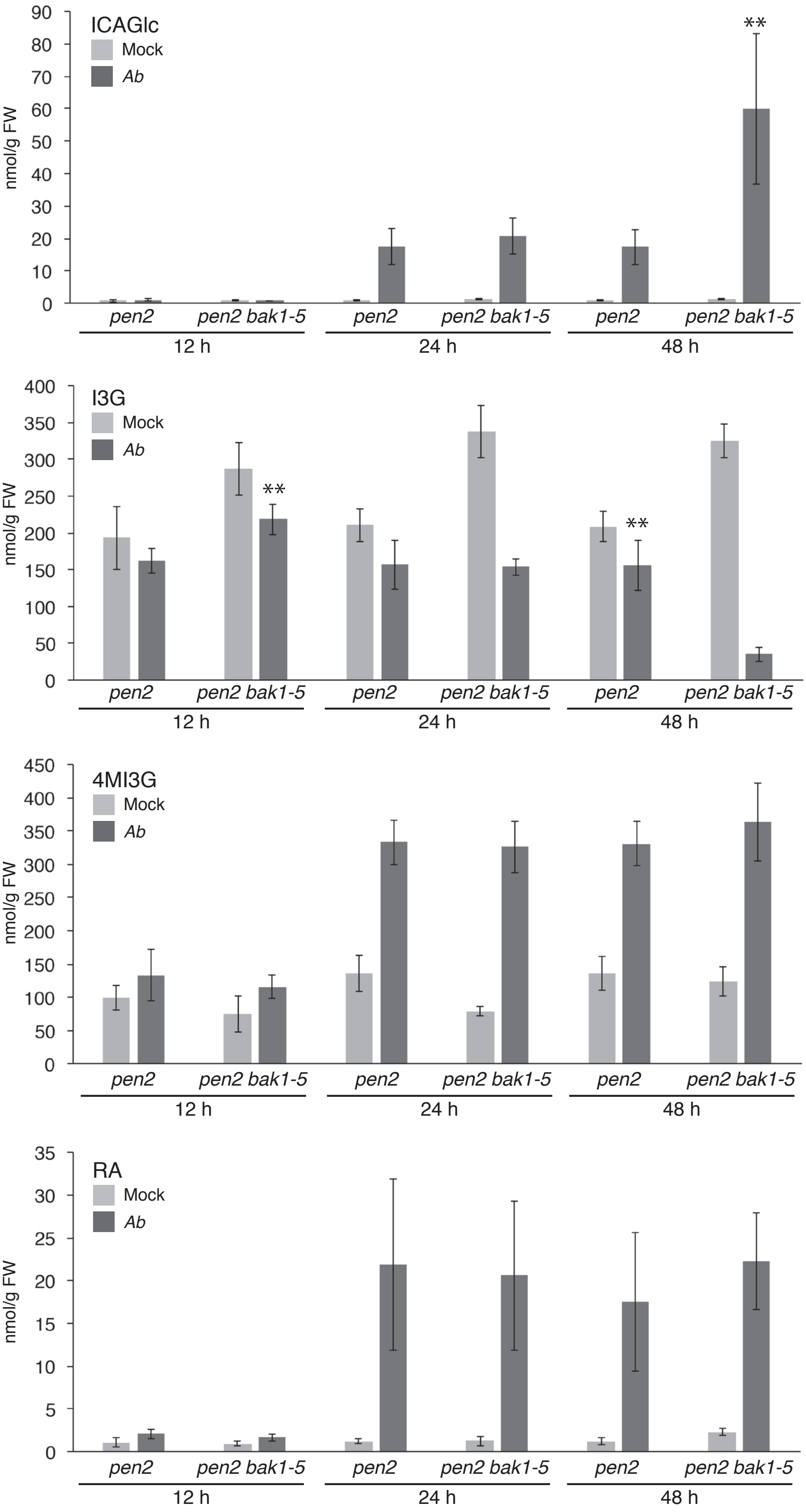


**Supplementary Fig. S7.** The analysis of ICAGlc, I3G, 4MI3G, and RA accumulation in both *pen2* and *pen2 bak1–5* mutants. Conidial suspensions (5 x $10^5$ conidia/mL) of *Ab* were spray-inoculated onto the tested plants. As a control, water was sprayed as a mock treatment. The samples were collected at 12, 24, and 48 hpi. Accumulations of ICAGlc, I3G, 4MI3G, and RA were determined. The means of metabolites (nmol/g FW) and SDs from four biological replicates are shown in the graph. Comparisons between *pen2* and *pen2 bak1–5* at each time point were conducted using two-tailed Student's *t* tests (**$P$ < 0.01).

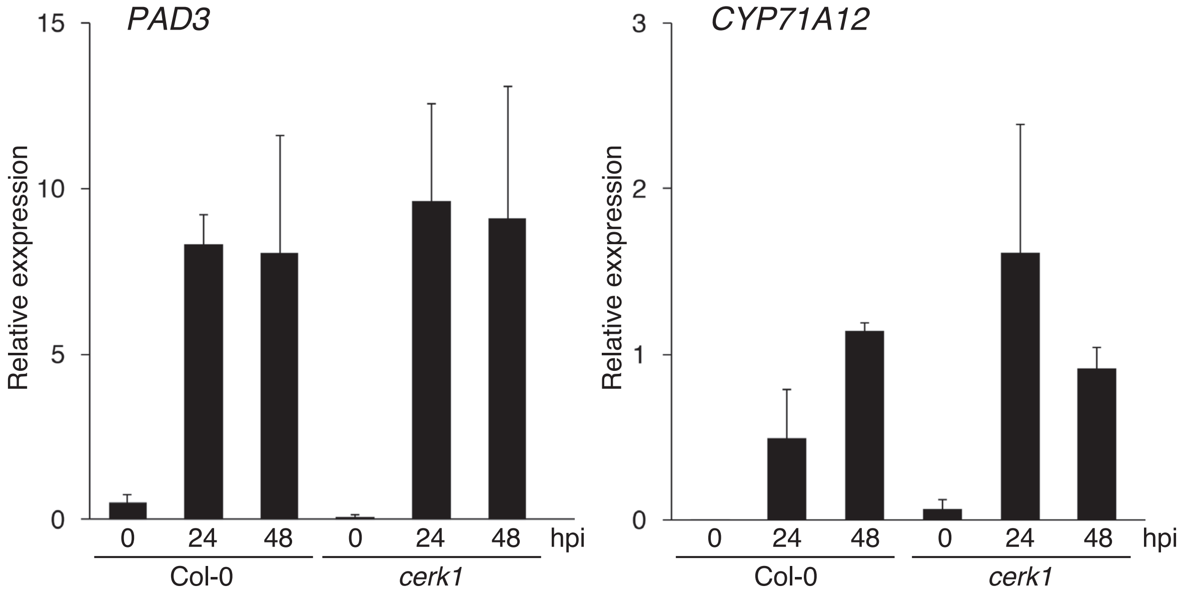


**Supplementary Fig. S8.** The expression of *CYP71A12* and *PAD3* triggered by *Ab* invasion is not dependent on CERK1. Conidial suspensions (5 x $10^5$ conidia/mL) of *Ab* were spray-inoculated onto 4–5-week-old plants. The samples were collected at 0, 24, and 48 hpi. Each gene transcript was quantified by RT–qPCR using the gene-specific primers listed in Supplementary Table S2. Values were normalized to the expression level of *UBC21*.

**Supplementary Table S1.** List of differentially regulated Arabidopsis genes in *pen2 bak1–5* plants after *Ab* invasion at 24 hpi

| Locus | Description | *pen2 bak1-5/pen2* [a] |
|---|---|---|
| Down-regulated in *pen2 bak1-5* | | |
| At5g52740 | copper transport family protein | 0.20 |
| At4g04500 | cysteine-rich receptor-like protein kinase 37 (*CRK37*) | 0.21 |
| At2g26400 | acireductone dioxygenase 3 (*ARD3*) | 0.21 |
| At4g23150 | cysteine-rich receptor-like protein kinase 7 (*CRK7*) | 0.29 |
| At3g57260 | beta 1,3-glucanase 2 (*BGL2*) | 0.30 |
| At5g55450 | bifunctional inhibitor/lipid-transfer protein/seed storage 2S albumin-like protein | 0.31 |
| At5g10760 | aspartyl protease family protein (*AED1*) | 0.33 |
| At2g14580 | pathogenesis-related protein 1 (*PRB1*) | 0.34 |
| At2g32680 | receptor like protein 23 (*RLP23*) | 0.36 |
| At5g40990 | GDSL lipase 1 (*GLIP1*) | 0.37 |
| At3g28510 | AAA-type ATPase family protein | 0.39 |
| Up-regulated in *pen2 bak1-5* | | |
| At1g29660 | GDSL esterase/lipase | 25.80 |
| At5g44420 | ethylene- and jasmonate-responsive plant defensin (*PDF1.2*) | 4.39 |
| At1g79400 | cation/H(+) antiporter 2 (*CHX2*) | 2.58 |

[a]Up-regulated genes were determined by a greater than 2.5-fold change of normalized signals in their expresion ratio (*pen2 bak1-5* vs *pen2*).
Down-regulated genes were determined by a smaller than 0.4-fold change of normalized signals in their expresion ratio (*pen2 bak1-5* vs *pen2*).
The values are the average ratios of four arrays.

**Supplementary Table S2**. List of primers used in this study

Primers used for RT-qPCR

| Gene name | Forward primer sequence (5'-3') | Reverse primer sequence (5'-3') |
|---|---|---|
| *CYP71A12* (AT2G30750) | CATTCCCTAAGCCTTCGGTAC | CTTGGAGTTTCTTCATAACA |
| *CYP82C2* (AT4G31970) | CATTTGGTTCGGGAAGAAGA | AGCCAGGGCTCTCAGTCATA |
| *PAD3* (AT3G26830) | TGCTCCCAAGACAGACAATG | GTTTTGGATCACGACCCATC |
| *UBC21* (AT5G25760) | CTGCGACTCAGGGAATCTTCTAA | TTGTGCCATTGAATTGAACCC |
| *AED1* (AT5G10760) | GCGTATACAGTATCGTTTACGGCG | GGTAGTGAGAGTTTGCCAGGGCC |
| *BGL2* (AT3G57260) | TGGATCACCGAGAAGGCCAGGG | GCCCACAAGTCTCTAAGGATTAG |
| *GLIP1* (AT5G40990) | GAGCTGATTTGGAGCGGACCTACC | CGAGTGATATATATCGCTCGCG |
| *RLP23* (AT2G32680) | GGAGTGGCTTGTCAAGATAATTGG | CCCAATTTTATCCTCATTTGCCCG |

Primers used for genotyping

| Mutant name | Forward primer sequence (5'-3') | Reverse primer sequence (5'-3') | LBa primer sequence (5'-3') | LBb primer sequence (5'-3') | Enzyme |
|---|---|---|---|---|---|
| *pen2-1* | TCAGGTAAATCAGTTCGAATCAAGAAC | TGAGGAAACCTGTTGGAGAAAGGATC | | | BamHI |
| *bak1-4* | ATTTTGCAGTTTTGCCAACAC | CATGACATCATCATCATTCGC | TGGTTCACGTAGTGGGCCATCG | | |
| *bak1-5* | AAGAGGGCTTGCGTATTTACATGATCAG | GAGGCGAGCAAGATCAAAAG | | | RsaI |
| *pepr1* | TCGTTTCGGATCACCTAATTG | TTTCACCTGTCAATCCGTTTC | TGGTTCACGTAGTGGGCCATCG | | |
| *pepr2* | TTGGATCCAACTCATTGAACG | ACGCCAGTCCATGTGAAATC | | GCGTGGACCGCTTGCTGCAACT | |

**Supplementary Table S3.** List of *Arabidopsis thaliana* lines used in this study

| *Arabidopsis thaliana* lines | Reference |
|---|---|
| Col-0 (WT) | |
| *pen2-1* | Lipka *et al.,* 2005 |
| *pen2-2* | Lipka *et al.,* 2005 |
| *pad3-1* | Glazebrook and Ausubel, 1994 |
| *cyp71A12* | Müller *et al.,* 2015 |
| *cyp71A12* | Rajniak *et al.,* 2015 |
| *bak1-4* | Chinchilla *et al.*, 2004 |
| *bak1-5* | Schwessinger *et al,* 2011 |
| *cerk1-2* | Miya *et al.,* 2007 |
| *pepr1 pepr2* | Yamaguchi *et al.,* 2010 |
| *cyp71A12 cyp71A13* | Müller *et al.,* 2015 |
| *cyp79B2 cyp79B3* | Zhao *et al.,* 2002 |
| *pen2 pad3* | Bednarek *et al.,* 2009 |
| *pen2 cyp82C2* | Pastorczyk *et al.,* 2019 |
| *pen2 cyp71A12* | Pastorczyk *et al.,* 2019 |
| *pen2 bak1-4* | In this study |
| *pen2 bak1-5* | In this study |
| *pen2 pepr1 pepr2* | In this study |
| *myb 34 myb51 myb122* | Frerigmann and Gigolashvili, 2014 |
| *pen2 cyp71A12 cyp71A13* | Pastorczyk *et al.,* 2019 |
| *cyp71A12 cyp71A13 myb 34 myb51 myb122* | Pastorczyk *et al.,* 2019 |
